# Nanoscale 3D DNA tracing in single human cells visualizes loop extrusion directly in situ

**DOI:** 10.1101/2021.04.12.439407

**Authors:** K.S. Beckwith, Ø. Ødegård-Fougner, N.R. Morero, C. Barton, F. Schueder, W. Tang, S. Alexander, J-M. Peters, R. Jungmann, E. Birney, J. Ellenberg

## Abstract

The spatial organization of the genome is essential for its functions, including gene expression, DNA replication and repair, as well as chromosome segregation. Biomolecular condensates and loop extrusion have been proposed as the principal driving forces that underlie the formation of chromatin compartments and topologically associating domains, respectively. However, whether the actual 3D-fold of DNA in single cells is consistent with these mechanisms has been difficult to address in situ. Here, we present LoopTrace, a workflow for nanoscale 3D imaging of the genome sequence in structurally well-preserved nuclei in single human cells. Tracing the in situ structure of DNA in thousands of individual cells reveals that genomic DNA folds as a flexible random coil in the absence of loop extruding enzymes such as Cohesin. In the presence of Cohesin and its boundary factor CTCF, reproducibly positioned loop structures dominate the folds, while Cohesin alone leads to randomly positioned loops. The 3D structure and size variability of DNA loops we observe in a large number of single cells allow us to formulate a data-constrained computational model of genomic DNA folding that explains how sparse and dynamic loops in single cells lead to the emergence of compact topological domains in averages of cell populations.

## Introduction

The spatial organization of the human genome covers a large hierarchy of spatial scales, from individual nucleosomes of a few nanometers to supra-chromosomal compartments of several microns, and plays a central role in regulating genome functions ^1^. Crosslinking and sequencing based chromosome conformation capture methods (e.g. 3C/Hi-C) have defined compartments at the megabase (Mb) scale and topologically-associating domains (TADs) of several hundred kilobases (kb) as conserved features of genome organization ^2,3^. While large scale compartments are believed to be driven by biological condensation ^4^, TADs have been shown to fall between CCCTC sequence elements bound by CTCF ^3^ and depend on the Cohesin complex, a molecular motor that can extrude DNA loops in vitro ^5–8^. Polymer simulations recapitulating HiC data have suggested that compact domains inside cells could thus arise from promiscuous loop extrusion in chromosomal DNA constrained by flanking CTCF sites ^9–11^. Recent observations of the mobility of fluorescently labelled CTCF-sites in live cells are consistent with a boundary function ^12^. However, direct evidence for the nanoscale structure of genomic DNA in single cells, that could address if individual loops are in fact present in human chromosomes and if their size, number and genomic position is consistent with our current models for genome architecture is still lacking.

Providing direct structural evidence for the nanoscale fold of genomic DNA in situ has been technically very challenging. Contact frequencies between genomic loci after crosslinking and sequencing as done in Hi-C do not directly measure the three-dimensional (3D) physical structure of genomic DNA ^13^ and attempts to reconstruct higher order DNA folds in cryo-preserved specimen by electron microscopy have failed due to the lack of sequence specific labels and the very high density of DNA in the nucleus ^14^. By contrast, light microscopy-based methods such as the multiplexed labelling of short unique genomic sequence elements by oligonucleotide fluorescence in situ hybridization (FISH) probes in principle allow direct, single-cell visualization of genome structure ^15–25^. So far however, comparatively harsh denaturing hybridization conditions were used that do not conserve the fragile nanoscale structure genomic DNA ^26,27^. Nevertheless, genome imaging could already reveal a strong cell-to-cell variability of the larger scale structural features seen by HiC ^15–25^.

In this study, we aimed to deploy the power of in situ genome imaging to investigate the nanoscale 3D folding of single chromosomal DNA molecules down to individual loops. We therefore developed a scalable high-resolution DNA tracing workflow that uses non-denaturing labelling with oligopaint FISH probes with a genomic resolution of better than 5 kb and preserves the nanoscale structure of the genome in a near native state. We combine this gentle, yet robust and precise labeling with rapid, automated sequential 3D imaging with a spatial precision of better than 25 nm. Using this novel workflow, which we term “LoopTrace”, we systematically investigate the 3D-folding of chromosomal DNA in multiple genomic regions from the kb to Mb-scale in single human cells. Our approach provides fast and reliable 3D tracing below the level of single DNA loops in thousands of individual cells.

By directly visualizing its 3D structure, we reveal that genomic DNA folds like a random-coil polymer in the absence of Cohesin. We furthermore show that individual Cohesin-dependent loops exist in only about 50% of the cells, and that these loops can only be reproducibly positioned if CTCF is also present. Finally, we also visualize the complex longer range DNA interactions of entire topological domains at the megabase scale in 3D. The size, frequency and genomic position of the DNA loops we measure directly in situ allow us to fully constrain a computational model of genome folding with experimental data. This data-driven model predicts that the cell can make more and longer DNA loops if needed and explains how sparse and dynamic loops in single cells cumulatively lead to the emergence of TAD-like features at the population level.

## Results

### LoopTrace enables nanoscale 3D DNA tracing with near-native structure preservation

To investigate which FISH protocol would best preserve genome structure and yet allow the high labelling efficiency required for reliable DNA tracing ^28^, we compared the previously used denaturing FISH approaches with enzymatic, non-denaturing FISH (CO-FISH/RASER-FISH; see Supplementary Note 1 and Supp. Fig. S1) ^29–31^. After systematic optimization, we found that non-denaturing FISH combines good hybridization efficiency of oligopaint probes to genomic DNA with near-native preservation of genome and nuclear structure, making it a suitable method to investigate the folding of chromosomal DNA at the nanoscale in situ in single human cells.

We next developed a sequential DNA-FISH imaging workflow, LoopTrace, to achieve reliable high-resolution 3D-DNA tracing at the scale of single loops. To establish a baseline, we first imaged the non-coding genomic region upstream of the MYC gene on the right arm of chromosome 8, which is predicted to represent open euchromatin (Supp. Fig. S2a) and contains no CTCF-binding sites (Fig. 1a). To resolve its structure, we targeted sets of oligopaint probes containing locus-specific barcodes to ten discrete 5-kb-loci spaced 5 kb apart. After the initial, gentle hybridization, the stably bound primary probes were then detected by sequentially binding and imaging unique readily exchangeable fluorescent 12-bp imager strands matching each locus barcode ^32^. We found that our workflow delivers a reliable and fully removable fluorescent label for each of the loci (Fig. 1b; Supp. Fig. S2b-d) that could be localized with better than 25 nm 3D precision (Supp. Fig. S2e-g). “LoopTrace” thus allowed reliable reconstruction of genomic DNA folds in 3D. The rapid imager exchange and imaging cycles allow a high degree of scalability and enabled measurements of more than one hundred genomic loci in ∼1000 cells in a typical overnight experiment.

**Fig. 1:**
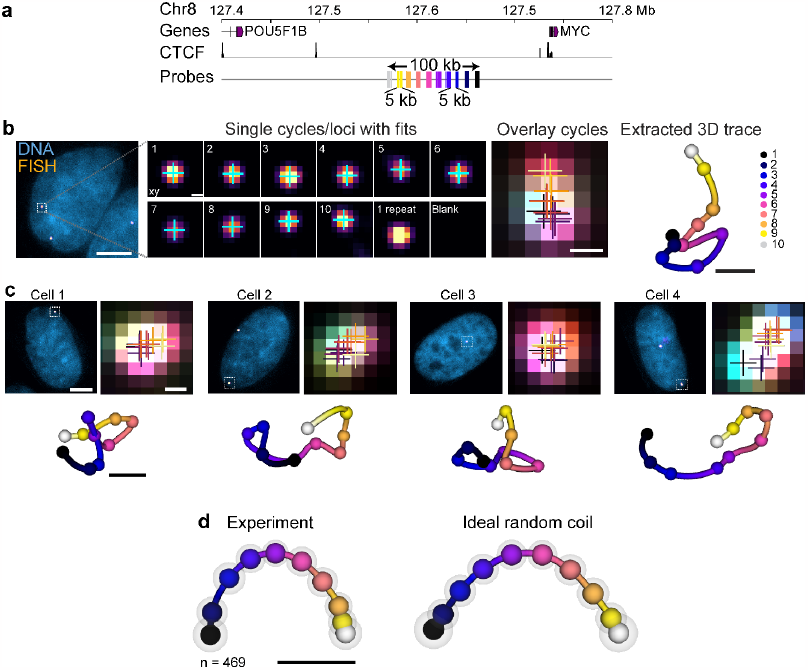
LoopTrace enables high precision 3D DNA tracing in non-denatured cells. **a**, Overview of a non-coding region upstream of MYC gene selected for DNA tracing pipeline validation with genes and CTCF sites (RPE Chip-seq^33^) highlighted. The probes cover 10 regions of 5 kb with 5 kb gaps, spanning a total of 100 kb. **b**, Examples of FISH signal, projection of sequential imaging steps and centroid fits, and resulting 3D DNA trace. **c**, Overlaid FISH signal with fits and resulting DNA traces from four representative cells. **d**, 3D consensus traces of experimental and simulated data calculated by general Procrustes analysis, with shaded spheres showing the standard deviation of each position. Data from n=469 traces from 3 independent experiments or simulated ideal random walks. Scale bars: 20 µm (overview), 5 µm (nuclei), 200 nm (FISH spots) and 100 nm (traces).

### A 100 kb open euchromatic region folds like a random coil

To get structural insights into the 100 kb euchromatic region upstream of the MYC gene, we compared the 3D traces between individual cells and found high cell-to-cell variability of DNA folding (Fig. 1c). Combining the 3D traces from hundreds of cells into a consensus trace by general Procrustes analysis or pairwise median distance map of all loci (see methods) showed no reproducible substructure (Fig. 1d, Supp. Fig. S2h,i). To test if our data can be explained by the behavior of DNA as a dynamic polymer, we set up a computational model for a random coil scaled to match our genomic target region. Both the consensus traces and pairwise distance maps showed very close agreement between experimental data and random coil model (Supp. Fig. 2i, r = 0.98). Interestingly, fitting the measured 3D distances to a power law gave a scaling exponent ν = 0.40 (Supp. Fig. S2j), significantly lower than the value of 0.5 expected for an ideal random coil or 0.588 for a highly solvated polymer ^34^, indicating that this 100 kb region without CTCF binding sites folds as a compacted random coil.

### LoopTrace visualizes single loops between convergent CTCF sites directly in 3D

To investigate if “LoopTrace” could resolve individual DNA loops in single cells, we next targeted 100-300 kb-scale genomic regions on four chromosomes predicted to have looping architectures due to the presence of convergent CTCF sites (Supp. Fig. S3a) ^3^. To resolve their 3D structure in situ for the first time, we tiled 25-30 FISH probes at 4 or 10 kb resolution and imaged them in diploid RPE-1 and the on average triploid ^35^ HeLa cancer cell line, respectively. Our data showed that the genome structure between these two human cell lines is conserved (Spearman’s r = 0.95±0.02), and at the population level consistent with publicly available Hi-C data (compare Supp. Fig. S3a, b and c), validating our technical approach.

To take full advantage of our data quality and gain a detailed and quantitative understanding of the physical 3D structure of a single DNA loop, we next imaged a 120 kb region with one pair of convergent CTCF sites near the MEOX1 gene on chromosome 17 with 10 kb genomic resolution (Fig. 2a). Consensus traces built from more than 2000 single cell folds clearly resolved loop features that brought the two CTCF sites into close proximity in 3D in about half of the individual RPE-1 and HeLa cells (Fig. 2b, Supp. Fig. S3c). The consensus fold of the MEOX1 region was thus clearly distinct from the random coil fold of the region devoid of CTCF sites near the MYC gene on chromosome 8 (compare Fig. 1d with Fig. 2b).

**Fig. 2:**
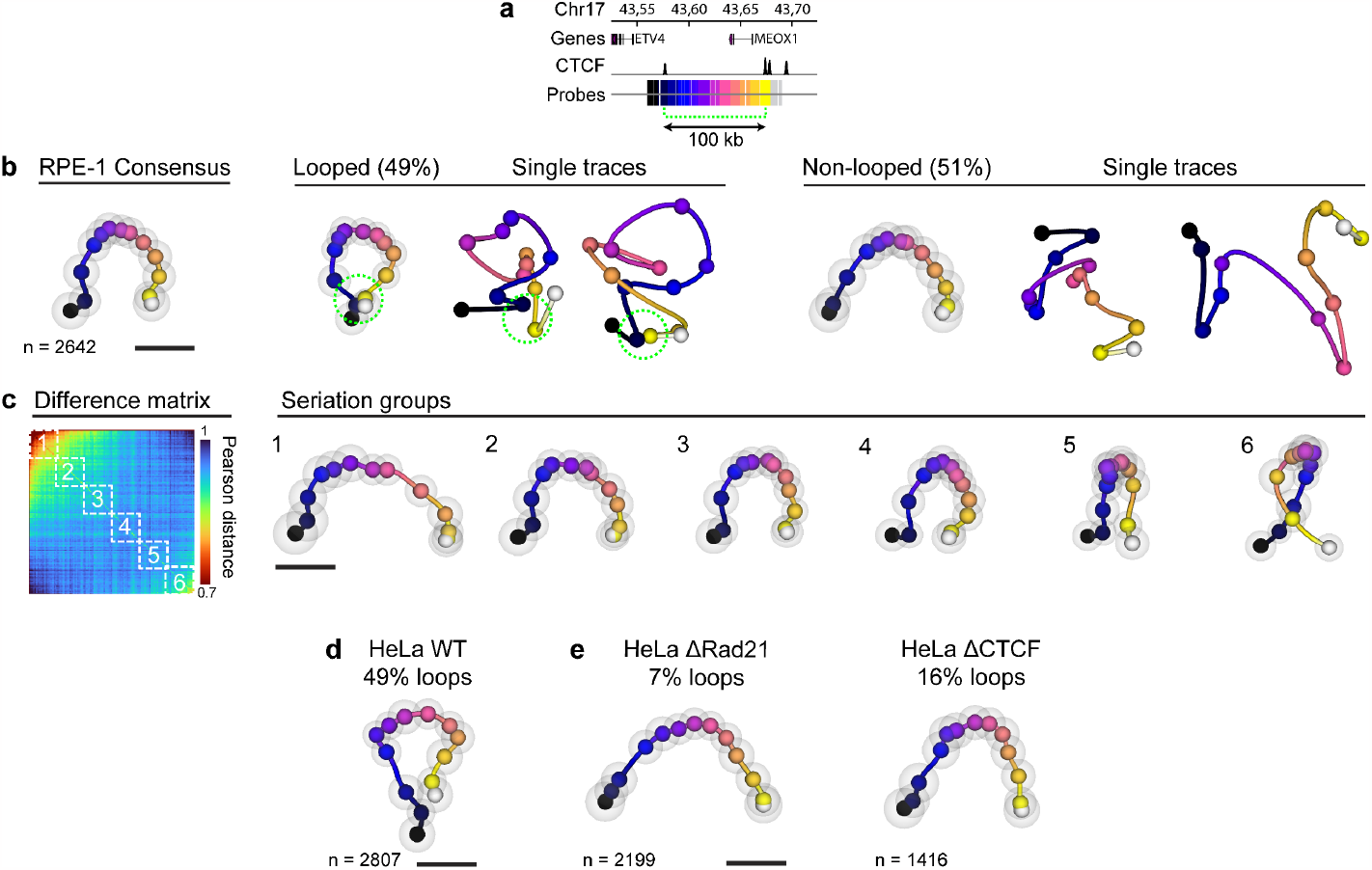
LoopTrace resolves single CTCF and Cohesin-anchored loops. **a**, Region targeted for tracing around the MEOX1 gene on chromosome 17. A pair of convergent CTCF sites predicted to form loops is indicated by the dashed line. **b**, RPE-1 consensus trace for the whole population, and consensus traces and single trace examples from the looping as well as non-looping subsets from the 120 kb MEOX1 region. Proximity (<120 nm) between the positions of the looping CTCF-sites indicated by dashed circle. **c**, Difference matrix 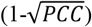 between single traces after sorting by the Fiedler vector (see methods), and the consensus traces from each numbered seriation group. n = 440 traces per group. **d**, Consensus trace from HeLa wild type (WT) cells. **e**, Consensus trace from HeLa Rad21-mEGFP-AID or HeLa CTCF-mEGFP-AID cells depleted of Cohesin or CTCF for 2h by auxin treatment. The number of traces for each experimental condition (2 replicates) are indicated. Scale bars 100 nm.

Comparing single cell loop traces showed a highly variable internal structure except for the proximity of the CTCF sites in looped traces (Fig. 2b) with 3D loop diameters that varied around 150 nm. Consistent with the MEOX1 loop, tracing a second 100 kb region on chromosome 9 with convergent CTCF sites at an even higher genomic resolution of 4 kb (Supp. Fig. S3d) also revealed clear loop structures with variable internal structure in about half the cells. Based on this data, we can conclude that single ∼100 kb sized CTCF-bounded DNA loops are present in about 50% of single human cells. These loops consistently exhibit close proximity to the loop anchors (CTCF sites) and high cell-to-cell heterogeneity of the structure inside the loop. The physical size of a 100 kb scale loop is on average 150 nm, highlighting that imaging and structure preservation at the nanoscale is necessary to visualize them.

### Unsupervised computational ordering of 3D folds suggests chromatin compaction is caused by loops

Our thousands of 3D traces from the same loop region represent snapshots that are likely to capture the dynamic genomic architecture of single cells in different states. To mine this rich data in an unbiased manner, we took an unsupervised computational approach to sort the traces into structurally similar groups to explore if we can detect transitions between different states. To this end, we ordered the traces from the MEOX1 loop region by spectral seriation (see methods). Visualizing six consecutive seriation groups as consensus traces revealed a progressive change from an open, random-coil like fold towards more and more compacted folds until the fully looped state was reached (Fig. 2c), very different from a simulated random coil that showed no such transitions (Supp. Fig. S4a). Seriation also revealed a similar progression from open folds to closed loops in the 3D traces of the second looping region on chromosome 9 (Supp. Fig. S4b).

Interestingly, although the overall DNA folds were similar, in HeLa cells the seriation analysis of the 120 kb MEOX1 region (Supp. Fig. S4c) also revealed differences in folding compared to RPE-1 cell line (compare Fig 2c, Supp. Fig. S3c). In particular, the seriation highlighted an additional interaction between the CTCF anchor at the MEOX1 promoter and regions inside the main loop, which may be related to the high level of MEOX1 transcription in HeLa cells ^36^ and/or potential enhancer elements in this region ^37^. Together, the unsupervised ordering into structurally similar groups of thousands of snapshots suggests that loops cause an increasing compaction of an initially open, loop-free genome architecture, consistent with progressive loop extrusion. Further, these results highlight the ability of our approach to detect distinct chromatin folding states between cells and cell types.

### Individual loops depend on Cohesin and their positioning on CTCF

If the looped folds we found in 50% of the cells indeed result from active extrusion, they should depend on the presence of Cohesin. To test this, we acutely depleted the Cohesin complex subunit Rad21 or CTCF in HeLa cells where these genes had been homozygously tagged with an AID degron tag^5^. Our 3D tracing data clearly showed both Cohesin and CTCF depletion led to an almost complete loss of loop-like compact folds (Fig. 2e; Supp. Fig. S3e), whose 3D traces revealed that the loop anchors were now physically far apart (Supp. Fig. S5a). In addition, the power law scaling exponent of all 3D distances increased significantly from v_control_ = 0.37±0.09 to v_DRad21_ = 0.47±0.03 after Cohesin depletion, very close to an ideal random coil of v = 0.5 (Supp. Fig. S5b). Seriation analysis further confirmed folds similar to simulated ideal random coils (Supp. Fig. S4c). We conclude that Cohesin is required for the formation of DNA loops and that its removal leads to an overall loss of compaction and 3D folds that are well explained by an unconstrained random coil up to a scale of 300 kb.

We next wanted to test the structural effect of the Cohesin positioning factor, CTCF. Rapid depletion of CTCF substantially reduced loop folds to below 20% of all cells in the 120 kb MEOX1 region (Fig. 2f, Supp. Fig. S5a). In contrast to Cohesin depletion, the traced regions in cells lacking CTCF remained overall similarly compact to control cells (Supp. Fig. S5b, v_DCTCF_ = 0.34±0.03 compared to v_control_ = 0.37±0.09). CTCF is thus required for reproducible positioning of loops, but not for compaction of the randomly folded genomic DNA. That higher compaction is maintained in the absence of CTCF could be due to continuous Cohesin loop extrusion without sequence specific constraints. This hypothesis was further supported by tracing a 300 kb region near the HIST1 gene cluster on chromosome 6 predicted to have only short range loops (Supp. Fig. S3a,b). Here, CTCF loss led to an increased frequency of long-range contacts (Supp. Fig. S3e; Supp. Fig. S5b), presumably due to Cohesin now being able to create longer loops without being stopped at closely positioned CTCF sites.

### The 3D structure of a TAD-scale genomic region reveals sparse and open loops

Given that we could capture different structural states consistent with underlying dynamic ^12,38^ transitions from random coil to loops at the ∼100-300 kb scale, we next explored if LoopTrace could also be used to resolve the architecture of larger, TAD-scale (1-2 Mb) genomic regions predicted to be more compact by biochemical approaches ^3^. To this end, we traced four 1.3 to 1.8 Mb sized regions on three chromosomes with typical TAD features in publicly available HiC data (Supp. Fig. S6a). We labelled these regions in HeLa cells with 10-12 probe sets spaced around 100-150 kb apart, decorating validated CTCF sites as well as intermediate loci to gain information on the behaviour of predicted loop anchors as well as internal loop structure (Fig. 3a; Supp. Fig. S6a).

**Fig. 3:**
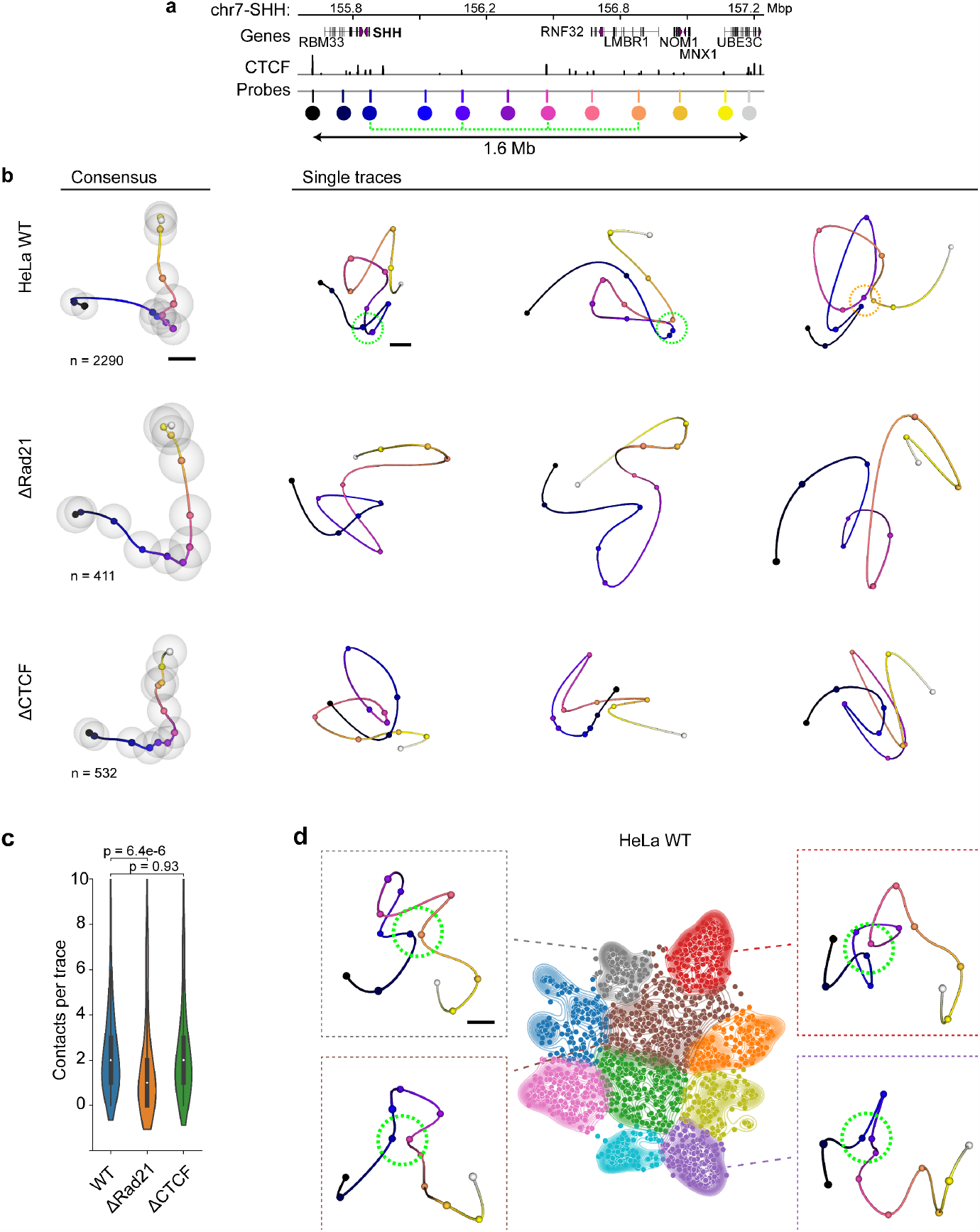
TAD-scale genomic DNA architecture in single cells. **A**, Overview of a genomic region (Chr7-SHH) targeted with 12 probes (highlighted by coloured circles) across 1.6 Mb. Each probe targets a ∼10 kb-sized region. Major CTCF sites predicted to form loops are indicated by dashed green line. **b**, Consensus traces and single trace examples from HeLa WT or HeLa Rad21-mEGFP-AID, CTCF-mEGFP-AID treated for 2 h with auxin. Scaling of the trace features is maintained from Fig. 2 but shown at lower magnification to cover the larger regions. Data from 3 (WT) or 2 (AID cells) replicates, number of traces per condition are indicated. Green dashed circles indicate <120 nm proximity between probe no. 3 at the TAD boundary and downstream CTCF sites, while cyan circles indicate multiple stacked contacts. **c**, Number of contacts (<120 nm) per trace in the Chr7-SHH region across the cell lines shown in (**b). d**, UMAP with clustering of single traces by a Bayesian GMM (see methods) from the chr7-SHH locus in HeLa WT cells. Median distance reconstructions of exemplary clusters and insets highlighting the internal TAD architecture (3^rd^ – 9^th^ probe positions, dominating loops highlighted with green dashed ring) as indicated by trace/cluster colour. n=2290 from 3 experiments, n=119-365 traces in each cluster. Scale bars are 100 nm.

3D DNA tracing of over 1000 single cells for each region revealed several close physical contacts and averaging their frequency within the cell population was again consistent with publicly available HiC data (Supp. Fig. S6a,b). Representing all data in 3D consensus traces revealed compaction inside the TAD regions as a conserved average structural feature in the four measured regions (Fig. 3b; Supp. Fig. S6c, d). However, interactively exploring the fold of individual 3D traces revealed an overall rather open architecture of the megabase sized regions with, on average, only ∼2 long-range physical contacts below 120 nm, typically between convergent CTCF sites (indicated by dashed circles in Fig. 3b,d; Supp. Fig. S6e). The average contact number was significantly reduced to below 1 upon Cohesin depletion, while CTCF depletion did not affect the number of contacts (Fig. 3c; Supp. Fig. S6c, e). Most long-range contacts (over 500 kb) resulted from stacking of two or more intermediate contacts (Supp. Fig. S6f; example in Fig. 3b indicated by orange circle), while single very large loops were observed only rarely.

To mine the wealth of TAD-scale 3D DNA folds from thousands of single cells in an unbiased manner, we next performed unsupervised clustering of all single traces using a Bayesian Gaussian mixture model (see methods) to group these structurally more complex 3D folds by similar structural features, using the Chr7-SHH locus as an example (Fig. 3d; Supp. Fig. S6g). Very interestingly, the 3D consensus traces of the clusters revealed that most automatically identified groups represented a specific physical contact between the upstream TAD boundary of the SHH gene and a different downstream loop anchor (Fig. 3a, d; Supp. Fig. S6g). In these clusters, the presence of these loops brought the boundary in close proximity with a loop anchor inside the TAD, while the lack of loops across the boundary to outside the TAD led to comparatively increased physical distances to these regions (Supp. Fig. S6g). These results provide a structural explanation for how a single strong anchor on one side of a loop can generate an effective boundary and yet lead to many different loop sizes and internal folds.

Together, these highly variable and open folds will average to a more compact domain at the population level (Fig. 3b). Overall, the 3D in situ structural data LoopTrace allowed us to generate at the larger scale of megabase sized genomic regions are fully consistent with our observations at the smaller, ∼100 kb, scale of single DNA loops. We can conclude that a typical TAD-scale region on average exhibits only sparse looping interactions inside a single cell, that these loops are rather open and of variable size but can be stacked to generate long-range physical proximity between genomically distant loci.

### A data-constrained polymer model predicts that loops and TADs are formed by a minor fraction of Cohesin complexes

The unsupervised clustering of Cohesin-dependent 3D folds into differently sized loops indicated that the loops we visualized are most likely generated by active extrusion in single cells. We therefore argued that we should be able to use our direct 3D structural constraints to set up a computer simulation that can predict how genomic DNA is folded in a single human cell assuming that it behaves as a random coil on which loops are progressively extruded. To this end, we adapted polymer/loop extrusion simulations that have been used to model genome structure ^9,10^, but constrained the parameters of our model systematically with the quantitative experimental data currently available for Cohesin complexes in single human cells (Supplemental Note 2; Fig. 4a). Our model includes the experimentally determined cellular protein concentrations ^39^, chromosome residence times ^39–41^, extrusion rate ^6,38^, as well as number and orientation of CTCF binding sites in the human genome ^3,39^. The main parameter we could not constrain was what fraction of the about 250 000 Cohesin complexes in a human cell is actively extruding loops.

**Fig. 4:**
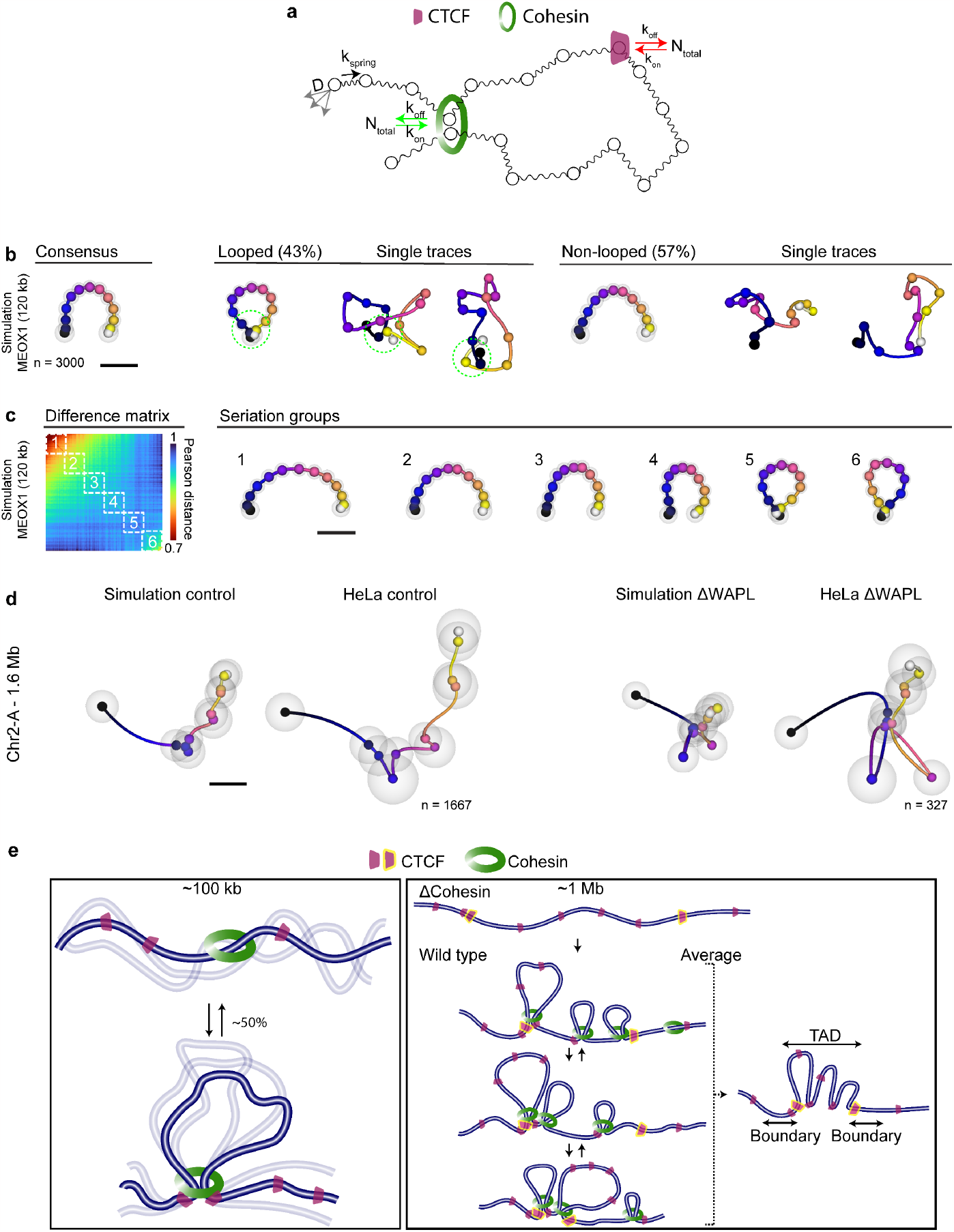
Data-constrained computational model of DNA loops and TADs. **a**, Schematic of the Rouse polymer/loop extrusion model which includes physiological values of the dynamic properties of chromatin, Cohesin and CTCF and is further constrained by 3D DNA tracing data. **b**, Consensus trace from simulations of the MEOX1 region on chromosome 9, together with looped and non-looped subsets and single trace examples, corresponding to the experimental data (Fig. 2b, c). **c**, Sorted matrix of Pearson’s distances between all traces and corresponding seriation analysis of the region shown in (**b). d**, Simulated and experimental consensus traces from a 1.6 Mb region on chromosome 2 traced with 11 probes with ∼150 kb spacing (see Supp. Fig. S6) in control cells and simulations. WAPL was depleted by auxin treatment of HeLa WAPL-Halo-AID cells for 2 h, in simulations Cohesin residence time was increased 20-fold^42^. **e**, Proposed experimentally parametrised loop extrusion model for chromatin architecture at the scale of 100 kb and 1 Mb.

To initially test the validity of our model, we set this fraction to 0% active Cohesin to reproduce our 3D DNA traces after Cohesin depletion as a reference for the random coil polymer and indeed obtained a very high accuracy fit (Supp. Fig. S7a, r^2^=0.95±0.08). To estimate the active Cohesin fraction, we then fit the 3D DNA traces from HeLa cells depleted of CTCF as well as control HeLa and RPE-1 cells. Strikingly, the model could only fit our single cell structural data with high accuracy (Supp. Fig. S7b-d, r_RPE-1_ = 0.92±0.05, r_HeLa_ = 0.88±0.06, r^2^_ΔCTCF_ = 0.91±0.07), when the fraction of active Cohesins was set to 20% and the abundance of CTCFs was set to the upper bound of the measured values ^39^. At this value, the model quantitatively reproduced the sparse abundance of loops, as well as their variable internal structures, leading to strikingly similar consensus traces, individual loops and progressive looping structures as those observed in our experimental 3D traces (compare Fig. 2b-d, experimental traces; Fig 4b, c, simulated traces). The simulation furthermore predicts that ∼100 kb sized loops are created by a single Cohesin complex while longer range contacts would result from the action of multiple Cohesins (Supp. Fig. S7c,d), providing an explanation for the stacked loops we found in our traces.

### As predicted by the model, increasing Cohesin binding to DNA further extends loops

Given the ability of the model to predict our experimentally observed 3D folds with a surprising level of accuracy and structural detail, we asked the model to make a testable prediction. As it is experimentally difficult to control the fraction of active Cohesin, we used the model to predict what structures would form if we forced active Cohesin complexes to bind 20 times longer to DNA, a situation that can be experimentally generated by removing its regulator WAPL^42^. In this in silico scenario, the model predicted an increased average length of non-stacked loops from 393±14 to 455±27 kb (Supp. Fig. S7e,f) and an increased proportion of stacked loops from 12±3% to 20±8%.

To validate this prediction experimentally, we acutely depleted WAPL from HeLa cells where the gene had been homozygously tagged with an AID degron tag^5^ and then retraced the 3D fold of this domain in over 300 individual cells. In striking agreement with the model prediction, we saw an increase in size and stacking of loops (Fig. 4d) that matched the simulated values with an error of less than 10% (Supp. Fig. S7f), suggesting that the parameters we use in the model are able to predict the 3D folding of genomic DNA with high accuracy up to the scale of 2 Mb.

Having validated the model placed us in a position to predict the key parameters of individual loops and TADs assuming the physiological situation, where all Cohesin regulators including WAPL are present (Fig. 4e). Here, our model predicts that in an average TAD-scale domain of around 1 Mb genomic DNA about four Cohesins interact with a dynamically exchanging pool of on average about 30 CTCF molecules to generate consistent loop contacts between the dominant CTCF boundary sites and internal anchors up to genomic distances of ∼300 kb. For a 100 kb single loop domain, the model would predict that 1 Cohesin interacts with 4 dynamically exchanging CTCFs leading to a single loop contact between convergent CTCF sites in about half of the cells (Fig. 4e).

## Discussion

The “LoopTrace” workflow we introduce here provides a precise, robust and scalable approach to investigate the nanoscale folding principles of the genome in situ in structurally well-preserved nuclei of single human cells. Our method does not suffer from the micrometer scale structural artefacts of denaturation-based FISH methods and shows good conservation of TAD sized domains at the 100 nm scale. However, even non-denaturing FISH methods show mild alterations of nuclear structure and in the future even more native methods to reliably target sequence specific probes to genomic DNA in single cells can be developed.

Our 3D DNA tracing with up to 25 nm/5 kb resolution provides direct, in situ measured structural data on the fold of the genomic DNA, which is key to understanding the structure function relationship of the genome. Based on indirect HiC data and computer simulations, Cohesin mediated loop extrusion had been proposed to be the mechanism that drives TAD formation ^9,10,38^. However, a direct visualization of the resulting looping conformations of genomic DNA has so far been lacking. Our 3D tracing data in single cells now provides such direct evidence. It shows that in the absence of Cohesin, genomic DNA at the scale of loops and TADs folds like a non-interacting random coil polymer. Onto this rather open and dynamic structure, loops are superimposed that depend on Cohesin activity and are positioned at specific genomic sites dependent on CTCF. Our quantitative structural data allowed us to construct a fully parametrized computer simulation, that predicts that single cells only use about 20% of all Cohesin complexes for extrusion of loops larger than ∼10 kb. Interestingly, this corresponds to the relative abundance of the Cohesin-STAG1 isoform, which could thus be sufficient to create the measured structures at the 100-2000 kb scale, which is in line with biochemical observations ^40,43^, and could suggest alternative functions for the STAG2 isoform of Cohesin ^40,43^. We note, that our model tends to predict a higher overall compaction of megabase scale regions than we measured experimentally (Fig. 4d; Supp. Fig S6b; Supp. Fig. S7e), which could indicate that the in vivo processivity of Cohesin is lower than measured in vitro ^6,38^.

We find that short-range (∼100 kb) loops between pairs of convergent CTCF sites are present in about 50% of cells, consistent with estimates from single-cell HiC ^44^ and live cell imaging ^45^, and our model predicts that they are typically formed by a single Cohesin complex. Longer-range contacts are sparser, around 1 per Mb region and result from stacking of two or three intermediate loops (Fig. 4e). The individual folds at this scale show different loop sizes and overall a rather open architecture. The complete set of TAD features, i.e. strong boundaries and high frequency of internal interactions thus does not exist in any single cell at any time point. Rather, TAD boundaries arise from the cumulative effect of dynamically positioned sparse looping interactions inside the extrusion range of strong CTCF sites in cell population data.

We anticipate that the LoopTrace approach we developed here, that combines near-native oligopaint FISH with scalable high-precision 3D DNA tracing, will be a valuable tool to directly investigate the structure-function relationship of the genome at the nanoscale in single cells. It is in principle scalable to whole chromosomes and even the whole genome ^28^, and in the future should enable a deeper understanding of the link between genome architecture and the functional state of individual cells in health and disease.

## Acknowledgements

We thank Merle Hantsche-Grininger for developing the electroporation protocol for replication domain labelling and critical reading of the manuscript; Franziska Kundel, Jonas Ries and Virginie Uhlmann for discussions; Magda Bienko and Jill Brown for sharing of FISH protocols; the EMBL Advanced Light Microscopy Facility (ALMF) for microscope support; Christian Tischer and Jean-Karim Hériché for help with data/metadata handling, and the electronic workshop and mechanical workshop at EMBL for assistance with the microfluidics setup.

## Author contributions

Conceptualization: JE and EB

Methodology: ØØF, KSB, NRM, CB, FS, TW

Software: KSB, CB

Validation: KSB, NRM, TW

Formal analysis: KSB

Investigation: ØØF, KSB, NRM

Data Curation: KSB, CB

Writing – original draft preparation: ØØF, KSB, JE

Writing – review and editing: All authors.

Visualization: KSB

Supervision: SA, JMP, EB, RJ, JE

Funding acquisition: KSB, SA, JMP, EB, RJ, JE

## Data and materials availability

Source image data and processed data from this study are available on the EMBL EBI Bioimage Archive/Biostudies at the following website: https://www.ebi.ac.uk/biostudies/studies/S-BIAD59. Publicly available datasets used in this study were GSE71831 (RPE-1 HiC) ^46^, GSE63525 (HeLa HiC) ^3^, ENCSR000DVI (RPE CTCF CHIP-seq), ENCODE: ENCSR000EDE (HeLa RAD21 CHIP-seq) and ENCODE ENCSR000AOA (HeLa CTCF CHIP-seq). Custom code for image acquisition and microfluidic control is available online at git.embl.de/grp-ellenberg/tracebot. Image analysis and LoopTrace analysis software is available online at git.embl.de/grp-ellenberg/looptrace. The primary oligo FISH probe design software is available at gitlab.com/Chromotrace_utils/probe-design.

## Materials and Methods

### Generation of HeLa cell lines

All AID-tagged HeLa cell lines used in this study were generated by homology-directed repair using CRISPR Cas9 (D10A) paired nickase ^47^. HeLa-SCC1-mEGFP-AID and HeLa-CTCF-mEGFP-AID were described before ^5^. Based on the cell line SCC1-GFP ^5^, we introduced a Halo-AID tag to the N-terminus of WAPL, generating Halo-AID-WAPL/SCC1-GFP. Subsequently, Tir1 expression was introduced by transducing a homozygous cell clone with lentiviruses using pRRL containing the constitutive promotor from spleen focus forming virus (SFFV) followed by Oryza sativa Tir1-3xMyc-T2A-Puro ^5^. The gRNAs sequences used were: CACCGCTAAGGGTAGTCCGTTTGT and CACCGTGGGGAGAGACCACATTTA. The primers used for genotyping were: TGATTTTTCATTCCTTAGGCCCTTG and TACAAGTTGATACTGGCCCCAA.

### Cell culture

RPE-1 cells (hTERT-immortalized retinal pigment epithelial cell line, ATCC No. CRL-4000, RRID: CVCL_4388) were grown in Dulbecco’s Modified Eagle Medium (DMEM/ F-12, Cat. No. 11320074, ThermoFisher) supplemented with 10% FBS (Cat.# 26140, ThermoFisher) and 1% Gibco Antibiotic-Antimycotic (Cat.# 15240096, ThermoFisher). HeLa Kyoto (HK) cells (RRID: CVCL_1922) were a kind gift from Pr Narumiya, Kyoto University. HK WT, HK Rad21-mEGFP-AID (+OsTir1), HK CTCF-mEGFP-AID (+OsTir1) and HK Rad21-mEGFP Halo-AID-WAPL (+OsTir1) were cultured in high glucose DMEM (Cat.# ;41965039, ThermoFisher) supplemented with 10% FBS, 1 mM sodium pyruvate (Cat.# 11360070, ThermoFisher), 2 mM L-glutamine (Cat.# G7513, Sigma) and 100 U/ml penicillin-streptomycin (Cat.# 15140122, ThermoFisher). Cells were incubated at 37 °C, 5% CO_2_ in a humidified incubator. At 70-80% confluence cells were trypsinized with 0.05% Trypsin/EDTA (Cat.# 25300054, ThermoFisher) and transferred to a new culture dish at appropriate dilutions every 2-3 days. For microslide seeding (µ-Slide VI 0.5 Glass Bottom, Cat.# 80607, Ibidi), trypsinized cells were counted and diluted to ∼5×10^5^ cells/ml in complete culture medium. For some experiments including AID-tagged cell lines, cell suspensions of each of the cell lines were separately labeled with either one or both of ViaFluor 488 SE and ViaFluor 405 SE fluorescent dyes (Biotium) according to the manufacturer’s instructions. Briefly, cell suspensions in 500 µL of PBS were mixed with the dye (1 µM) for 15 min at 37°C and quenched by addition of 500 µL of cell growth medium and incubation for 5 min. Cells were centrifuged and resuspended in 500 µL of culture medium, incubated for 25 min at 37°C and finally centrifuged again and resuspended in fresh medium at 5×10^5^ cells/ml. Labeled cells were mixed in equal amounts, seeded in Ibidi dishes and grown for 24 h in the presence of 40 µM BrdU: BrdC mix (3:1) and 200 uM auxinole (Hycultec) to inhibit background degradation ^48^.

To deplete AID-tagged proteins, auxinole was washed out 22 h after cell seeding, and replaced with 500 µM Inole-3-acetic acid (IAA, Sigma) for 2h. Degradation of AID-tagged proteins in HK cells under these conditions was previously determined ^5^, and verified again in this work by incubating HK Rad21-mEGFP-AID cells with IAA and/or auxinole for different times (from 15 min to 4 h) and quantifying the amount of target protein left by capillary electrophoresis (Jess Simple Western, ProteinSimple) (Supp. Fig. S8a). Protein normalization was achieved by determining the total amount of protein loaded in each capillary with a fluorescent dye that binds to all amino groups in proteins (PN reagent, ProteinSimple).

### Primary FISH probe design

Unique FISH probes were designed using the human reference genome GRCh38.p12. Sequences used are listed in Supplementary Table 1. For the probes targeting the MYC locus, we used the ChromoTrace probe designer utility (see code availability). Briefly, sequences were excluded if they were highly repetitive, if they had high sequence similarity elsewhere in the genome and if their GC content was under 30% or over 70%. To test for any off-target binding sites of the remaining probes the genome was searched for areas which have an exact match of 7 nt or more with a probe. These possible off-target matches were then filtered using a sensitive filtering scheme based on spaced seeds to remove lower similarity matches. For the remaining off-target matches, the melting temperature was estimated using the algorithm presented in ^49^. Probes for which the highest off-target melting-temperature was below 58 °C (calculated at 50mM [Na^+^]) were selected until the desired number of probes in a region was achieved. This pre-computed probe library could be queried in a fast and flexible way to design FISH probes targeting any locus. Docking sequences for secondary imagers were appended (see below), and probes were ordered as pooled libraries (IDT, oPools). For the probes targeting regions on chromosomes 2, 4 and 6, 7, 9, 12 and 17, we used OligoMiner ^50^ with genomic target length between 36 and 42 nt, T_m_ between 42 and 50°C (accounting for 2XSSC with 50% formamide), and a minimum 2 nt spacing. Picked probes were filtered by the 42 °C LDA model with 0.8 stringency and a maximum of 5 or 10 off-target 18 nt kmers. Up to 100 probes in 10 kb regions were used per position. Docking sequences for secondary imagers (two identical barcodes for each position and one for each region per oligo) and amplification primers were appended, and probe libraries were ordered from Genscript (Precise Synthetic Oligo Pools) and amplified (see below).

### Imager and primer sequence design and labeling

The secondary imaging probes matching the 100 kb-scale MYC library were designed using NUPACK ^51^. Here, we aimed for a free energy of ∼-17 kcal/mol at an ion concentration of 50 mM NaCl, 10 mM MgCl_2_ and a temperature of 24 °C. Inspired by the sequences used in ^32^, this free energy allows stable hybridization during image acquisition and, at the same time, efficient removal of the imaging probes with the stripping buffer. To ensure orthogonality between different sequences (no cross-reactivity), we checked that the free energy of a sequence to every other sequence was larger than -10 kcal/mol. Atto-565 labelled imagers (D1-10) were purchased from MWG Eurofins.

Further extending the number of imager sequences, 12-mer imagers targeting the probes for regions on chromosomes 2, 4, 6, 7, 9, 12 (named Dp1-38, 101-114) were sourced from a list of 12 bp human nullomers ^52^, filtered for low secondary structure (ΔG > -0.2 kcal/mol using Primer3 ^53^), T_m_< 40°C (at ion concentration 390 mM), GC-content 40-50%, less than 3 consecutive identical nucleotides, and few high temperature (> 28 °C at 390 mM) heterodimers in the set. 11 and 10 bp off-target sites for the 12 bp probes were detected by BLAST+ (blastn-short, E-value < 10000), and the probes were sorted ascendingly for 11 bp off-targets. 12-mer oligo imagers were ordered with 3’ or 5’-azide functionality (Metabion), reacted with Atto647N-alkyne or Atto643-alkyne (Attotec) using click chemistry (ClickTech Oligo Link Kit, Baseclick GmbH) according to the manufacturer’s instructions with 100 µM oligo and 2-fold excess dye. Labeled oligos were purified by diluting to 20 µM in TE buffer, then adding 5-fold excess water-saturated n-butanol (Sigma), vortexing for 10 s, centrifuging (14 000 g, 1 min), extracted once more, and carefully pipetted from underneath the n-butanol layer into a fresh tube.

PCR primer sequences were selected from 20-mer sub-sequences of orthogonal 25-mer sequences ^54^, filtered for 55 °C > T_m_ > 60 °C (at 50mM, primer3), less than 4 identical nucleotide repeats, and starting and ending with a C/G. The primer sequences were further screened for compatibility with the 12-mer docking sequences by ensuring heterodimer T_m_ under 15 °C (primer3, 390 mM ions) and under 40 °C hairpin T_m_ (primer3, 390mM ions) of sequences composed by combining all possible pairs of 10 000 filtered 20-mers with the T7 in vitro transcription promoter and 200 12-mers selected above. Forward and reverse primers were paired and sorted by the number of 12-mers the set was compatible with. All oligo sequences used in this work are listed in Supplementary Data 1.

### Amplification of FISH Libraries

The FISH library targeting chromosomes 2, 4, 6, 7, 9 and 12 was amplified from synthetic oligonucleotide pools (Genscript), through PCR amplification followed by in vitro transcription, reverse transcription and ssDNA purification, as described before ^55,56^.

Briefly, the PCR amplification protocol for each library and primer pair was optimized by monitoring the progression of the reaction in real time with a qPCR machine to observe the minimum number of PCR cycles necessary to reach the amplification plateau. PCR was performed using 75 µl 2x Phusion High-Fidelity PCR Master Mix (ThermoFisher), 15 ng oligonucleotide pool, 0.5 µM of each primer and water up to 150 µl. The following PCR protocol was used: (i) 98°C for 3 min, (ii) 98°C for 10 s, (iii) 66°C for 10 s, (iv) 72°C for 15 s. Steps (ii) to (iv) were cycled until the reaction approached its amplification plateau (14-18 cycles). The amplified dsDNA template was then purified with a DNA Clean & Concentrator-25 kit (Zymo Research) and eluted in 50 µl of water. 5 µl of this sample was set-aside for quality control by gel electrophoresis.

In vitro transcription was done using the HiScribe T7 Quick High Yield RNA Synthesis Kit (NEB), following the manufacturer’s instructions for short transcripts. Reaction was set up as follows: 1.5 µg DNA template, 10 mM NTP buffer mix, 6.25 µl T7 RNA Polymerase mix, 250 U RNAsin Plus RNAse inhibitor (Promega) and water to 160 µl. Amplification proceeded for 16 h at 37°C. At this point, 1 µl of this sample was set-aside for quality control.

ssDNA amplification from template RNA was performed by reverse transcription, using: 150 µl of unpurified in vitro transcription reaction, 1.7 mM dNTP mix, 19 µM forward primer, 240 U RNAsin Plus RNAse inhibitor (Promega), 1,200 U Maxima H Minus Reverse Transcriptase (ThermoFisher), 60 µl of 5x Maxima Buffer and water up to 300 µl. The reaction mix was incubated for 1h in a water bath at 50°C. Template RNA was subsequently degraded by addition of 150 µl 0.5M EDTA and 150 µl 1N NaOH, and incubation for 15 min in a water bath at 95°C.

Amplified ssDNA library was purified with the HiScribe T7 Quick High Yield RNA Synthesis Kit (NEB). For this, the reaction solution (600 µl) was mixed with 1.2 ml Oligo binding buffer and 4.8 ml 100% ethanol, and then loaded into 2x 100-μg capacity purification columns. Columns were washed twice with 250 µl washing buffer and ssDNA was eluted with 100 µl nuclease-free water per column. Library concentration was measured by spectrophotometry in a Nanodrop.

Finally, to assess the specificity and efficiency of the amplification reaction, the intermediate products of each amplification step (PCR-amplified dsDNA, RNA and final ssDNA) were analyzed by Gel electrophoresis in denaturing conditions (Novex TBE-Urea Gels, 15%, ThermoFisher).

### Non-denaturing FISH

Non-denaturing FISH (RASER-FISH) was adapted from previously described protocols ^30,31^ to be performed in 6-channel microslides (µ-Slide VI 0.5 Glass Bottom, Ibidi), while minimizing DNA denaturation and genome structure perturbations and optimizing FISH signal for DNA tracing. Briefly, cells were seeded in microslide channels at a final concentration of ∼5×10^5^ cells/ml in 120 µL cell culture media containing 40 μM BrdU:BrdC mix (3:1, Cat.# B5002, Sigma and Cat.# 284555, Santa Cruz Biotec) and grown for 17-24 h. Afterwards, cells were washed once with PBS and fixed with 4% PFA (v/v, Cat.# 15710, EMS) in PBS for 15 min. Free aldehyde groups were quenched with 100 mM NH_4_Cl (Cat.# 213330, Sigma) in PBS for 10 min and cells were permeabilised in 0.5% Triton X-100 (v/v, Cat.# T8787, Sigma) in PBS for 20 min. To further sensitize BrdU/C-labelled DNA to UV light, cells were treated with 0.5 μg/ml DAPI (Cat.# D9542, Sigma) in PBS for 15 min and washed once in PBS. Microslides were exposed to 254 nm UV light for 15 minutes in a Stratalinker 2400 UV crosslinker, and cells were finally treated with 1U/μL Exonuclease III (Cat.# M0206, NEB) in 1X NEBuffer 1 at 37 °C for 15 min in a humid container. All incubations were performed at room temperature, unless stated differently. For primary probe hybridization, cells were first incubated at 37 °C for 1 h in buffer H1 (50% formamide (FA, Cat. # AM9342, ThermoFisher), 10% (w/v) dextran sulfate (D8906, Sigma) (Cat.# R6513, Sigma) in 2xSSC (Cat.# AM9763, ThermoFisher) and afterwards with primary probes (∼2nM per primary probe) diluted in H1. Primary probes were hybridized overnight at 42°C in a humid chamber. After primary hybridization, cells were washed twice in 50% FA in 2xSSC buffer at 35°C (7-10°C below probe on-target T_m_) for 5 min, followed by 3 washes with 2xSSC containing 0.2% (v/v) Tween-20 (Cat.# P9416, Sigma). For the probe library targeting regions on chromosomes 2, 4, 6, 7, 9, 12 and 17, RNAse A was not included in the hybridization buffer and was replaced by an RNAse H (NEB) treatment (1:100 in RNAse H reaction buffer for 20 min at 37°C) after primary hybridization to reduce non-specific background. For non-sequential secondary probe hybridization, fluorescently labelled 20-mer imagers were diluted to a final concentration of 20 nM in buffer H2 (25% FA, 10% (w/v) dextran sulfate, 0.1% (v/v) Tween-20, 2xSSC buffer) and incubated for 2 h at 30 °C. Washing steps were then conducted with 25% FA in 2xSSC buffer for 5 min, followed by washes with 2xSSC containing 0.2% (v/v) Tween-20. Before imaging, DNA was stained with 0.2 μg/ml DAPI for 10 min.

### Classical FISH

Standard FISH protocols including heat denaturation of DNA were performed in parallel to compare with the non-denaturing FISH procedures. RPE-1 cells were seeded in microslides, fixed and permeabilized as above and treated with 0.1M HCl (Cat.# 320331, Sigma) for 15 minutes. For primary probe hybridization, cells were incubated for 1h in buffer H1 at 37 °C, followed by 1h incubation in the same buffer containing the primary probes and a 3 min heat incubation step in a ThermoBrite slide hybridization system (Leica Biosystems), either at 75°C or 86°C. The temperature during incubation was verified with an additional external digital thermometer. Hybridization was then continued overnight at 42 °C. Washes and secondary hybridization steps were as described for the non-denaturing FISH protocol.

### Pulse DNA labelling with fluorescent nucleotides

Fluorescent labelling of TAD domains was adapted from ^57^ with some modifications. Briefly, cells were subjected to cell cycle arrest with 1 µg/ml aphidicolin (Cat.# A0781, Sigma) for 9 h, and subsequently allowed to recover and enter S-phase by incubation in fresh DMEM media for 20 min. Afterwards, cells were trypsinized, washed with PBS and resuspended in Resuspension Buffer R (Neon™ Transfection System 10 µL Kit, Cat.# MPK1025, Invitrogen) at a final density of 5.5 × 10^6^ cells/ml. dUTP-AF647 (Cat.# NU-803-XX-AF647-L, Jena Biosciences) was added at a final concentration of 60 µM and cells were electroporated with the following pulse parameters: 1 pulse, 1100 v, 20 ms with the Neon transfection system (Invitrogen). Labeled cells were grown in complete DMEM media, passaged after 1 day and seeded in microslide chambers 3 days after transfection. For non-denaturing FISH experiments, 40 µM BrdU/C was added at the seeding step.

### SIM image acquisition and analysis

RPE-1 cells pulse-labeled with fluorescent nucleotides were incubated with 1µg/mL Hoechst 33258 (Cat.# 14530, Sigma) in 1xPBS for 30 min. To avoid photobleaching, samples were imaged in an oxygen scavenger buffer system (100 mM tris, 50 mM NaCl, 0.5 mg/mL glucose oxidase, 40 μg/mL catalase, 10% glucose, 1 mM trolox for imaging before FISH, with tris and NaCl replaced with 2XSSC for imaging after FISH). SIM images were acquired on a Zeiss Elyra 7 with a 63X 1.4NA oil immersion objective using lattice SIM mode. Raw pixel size was 97 nm (1280 × 1280 pixels), z-spacing was 144 nm and camera (pco edge) exposure time was 50 ms. DNA (Hoechst, 405 nm laser) and replication domain (AF647, 642 nm laser) channels were acquired sequentially. Raw SIM images were processed in Zen Black 3.0 using standard settings except sharpening was set to “weak”, resulting in images with twofold increase in pixels in each dimension. To compare images acquired before and after steps in the FISH protocol a global 3D translational drift correction was first applied using SciPy ^58^. Single nuclei were detected and segmented using Cellpose ^59^, and the central plane of the nucleus was identified as the plane with the highest integrated intensity, and the corresponding plane was found in the comparison image as the plane with highest Pearson correlation. 2D slices of the central plane and the central plane + 8 slices were used for comparison. Maximum projections were used for the replication domains, and individual clusters of replication domains were segmented by Otsu thresholding. Images were cropped to the detected nuclei or replication domains, and a second pixel-level drift correction was applied to both rotation and translation (imreg-dft package). Finally, a subpixel translational registration was applied, before the pixel-wise Pearson correlation coefficient between the aligned images was calculated. The values for the two selected nuclear planes were averaged.

### Automated fluidics setup

A custom fluidics setup was built using a commercial 3-axis 30×18 cm GRBL controlled CNC stage. A syringe needle was mounted in place of the CNC drill head and connected using 1 mm i.d. PEEK and silicone tubing (VWR) to a male Luer adapter, which connected to the sample microslide. The microslide output tubing was connected to a mp6-liquid piezo micropump (Bartel’s mikroteknik), connected to an MP-X controller (Bartel’s mikroteknik) running at 130 Hz and 180 Vpp when dispensing, giving a flow rate of ∼3.5 mL/min, or a CPP1 peristaltic micropump (Jobst Technologies, Freiburg, Germany) running at full speed giving a flow rate of ∼1 mL/min. Probe, wash and imaging buffers were kept in 96 well or 4 well deep plates covered in parafilm (Cat.# P7793, Sigma) and placed on the CNC bed, and the appropriate solution was chosen by moving the syringe needle into the liquid through the parafilm. The automated control software was written in Python and was run on a PC connected to the MP-X controller/Jobst Pump Driver and GRBL CNC board. Detailed build and running instructions are described in the accompanying code repository (see code availability).

### 3D DNA trace acquisition

Cells on microslides were prepared for FISH as above, omitting secondary hybridization. 100 nm fluorescent fiducial beads (infrared Cat.# F8799, ThermoFisher or red Cat.# F8801, ThermoFisher) were diluted 1:20 000 in 2XSSC and applied to the sample for 10 minutes before washing in 2XSSC. 12-mer Atto565-labeled or Atto647N-labeled secondary imaging oligos were diluted to 20 nM in hybridization buffer (5% ethylene carbonate (EC, Cat.# E26258, Sigma) in 2XSSC/0.2% tween). Washing buffer was 10% FA in 2XSSC/0.2% tween, imaging buffer was 2XSSC with 1µg/mL Hoechst 33258 and probe stripping buffer was 30% FA in 2XSSC/0.2% tween. Probes were hybridized for 3 minutes and washed for 1 min before switching to imaging buffer and imaging. After imaging, probes were stripped for 2 minutes, then wash buffer was flowed through for 1 minute before the next round of probe hybridization. Imaging was performed on a Zeiss Elyra 7 widefield system using a 63X1.46NA oil immersion objective, a pco edge sCMOS camera and HILO mode, or a Nikon TI-E2 with a Lumencor Spectra III light engine, a 60X1.4NA oil immersion objective and an Orca Fusion CMOS camera and widefield mode. 3D stacks of Hoechst (405 nm laser), FISH probes (561 or 642 nm excitation) and fiducial beads (561 or 642 nm excitation) were acquired as sequential frames at each z-position. The pixel size was 97 nm (Zeiss Elyra) or 111 nm (Nikon Ti-E2), image size 1280×1280 (Zeiss Elyra) or 2304×2304 (Nikon Ti-E2) and z-spacing 150 or 200 nm, and the camera exposure time was 50 ms (Zeiss Elyra) or 100 ms (Nikon Ti-E2). 15-60 fields of view were acquired per round of hybridization. Total acquisition time for a typical experiment with 20-30 fields of view and 14-40 hybridizations was 4 - 20h. The Zeiss microscope was controlled using Zen 3.0 software (Carl Zeiss), with the MyPic macro ^60^, while the Nikon Ti-E2 was controlled using NIS Elements 5.2.02 (Nikon). For both systems custom Python software for automation and synchronization with the microfluidic software was used (see code availability). In experiments with multiple mixed ViaFluor labeled cell lines, the entire channel was imaged before FISH with 405 nm (VF405) and 475 nm (VF488) excitation and a 20X 0.8 air objective, relabeled with DAPI (100 ng/mL in PBS for 15 minutes) and imaged again using the same settings before proceeding to DNA tracing as above.

### DNA trace fitting

Custom python software based on SciPy 1.7.3 ^58^, pandas 1.4.4 ^61^, scikit-image 0.19.3 ^62^, scikit-learn 1.0 ^63^, pysimplegui, dask 2022.7.0 ^64^ and napari 0.4.16 ^65^ was developed to extract, fit and analyse DNA traces from the sequential FISH images (see code availability). Raw Zeiss CZI or Nikon ND2 images were read with czifile 2019.7.2 or nd2 0.4.4 python packages respectively, converted to the ZARR file format using (zarr 2.13.2) and deconvolved with a theoretical microscope-appropriate point spread function (10-30 iterations) or an experimental point spread function (60 iterations) with a Richardson-Lucy iterative algorithm (FlowDec 1.1.0 ^66^). Subpixel drift correction was calculated for all hybridization rounds by upsampled cross-correlation or Gaussian centroid fitting of 50-200 segmented fiducial beads and averaged. Regions of interest (ROIs) with FISH signal were automatically identified based on an empirical threshold in frames with imagers targeting most or all positions in a region for brighter signal. For weaker signal, enhancement of spot-like signals for detection using a difference of gaussian filter. When multiplexing tracing of multiple regions in the same cell, different regions were simultaneously traced and individual regions were identified by hybridization with region-specific probes, and ROIs were detected for each region in the respective frames. Nuclei were detected using Cellpose and only ROIs inside nuclei were included. Detected ROIs were quality controlled by manual spot-checks. For decoding VF405 and VF488-labeled cell lines (see Supp. Fig. S8a), nuclei images were cross-correlated with the low magnification images acquired before FISH treatment, then VF405 and VF488 intensities in 5-pixel dilated nuclear masks were manually selected on a scatterplot to decode the cell line identity.

The ROIs with FISH signals were cropped and the 3D image from each hybridization was fit to a 3D Gaussian function by least squares minimization, with the centre defining the coordinate in the corresponding DNA trace.

### DNA trace analysis

3D DNA traces were analysed using SciPy 1.7.3 ^58^, scikit-learn 1.0 ^63^ and umap-learn 0.5.3 ^67^python packages and visualized using Seaborn 0.11.2 ^68^. All fits were quality controlled by cut-offs on signal to background, fit standard deviation and distance to the multiple-probe/regional probe signal to avoid fitting of spurious background signals. Due to the non-denaturing FISH procedure, no traces from sister chromatids are generated and so do not need additional filtering. DNA traces were filtered for minimum percentage of quality-controlled fit loci of 67% for consensus traces, distance/contact maps and pairwise similarity measures, and 80% for contact counting.

Pairwise distance maps and contacts maps were calculated by directly calculating median pairwise distances from the measured 3D coordinates, and the frequencies of those distances less than a cut-off (set to 120 nm). Consensus traces were generated by a general Procrustes alignment using an iterative pairwise alignment to an initially randomly chosen template. Pairwise alignments were performed by 3D rigid alignment using singular value decomposition (SVD) without scaling to minimise the error (RMSE) ^69^. A quadratic spline interpolation was used to connect the fit loci coordinates for visualization. 3D traces were visualized with mayavi ^70^. Consensus traces of multiple traces were overlaid with the per-loci standard deviation of the single traces compared to the Procrustes consensus.

Seriation and clustering of traces was performed by calculating all pairwise distances of each trace, taking the reciprocal of each value to emphasise shorter distances, and then calculating the pairwise difference matrix between all traces with the Pearson distance 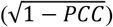 as the distance metric. Seriation was performed by ordering the traces according to the second largest eigenvector of the difference matrix (Fiedler vector) ^71^ and splitting the resulting ordered traces into six groups of equal size. Clustering of traces was done after UMAP dimensionality reduction (umap-learn) with n_neighbor = 20 and min_dist = 0 by Variational Bayesian estimation of a Gaussian mixture (scikit-learn) with n_components = 10 and other parameters at default values. To rule out that the obtained clusters could arise by chance, we compared average intra-cluster distances (0.697±0.11) to intra-cluster distances after randomly assigning traces to clusters of corresponding sizes (0.92±0.01), which validated that real clusters grouped significantly more similar traces than by random chance (Supp. Fig. 8b).

### Simulations

Ideal random coils were generated using 3D random walkers, where each step was in a random direction and had a length drawn from a normal distribution with a mean corresponding to the genomic length between each probe scaled by the empirical factor of 24.7 nm kb^-(1/2)^ found in our tracing data of the 100 kb region near the MYC gene, and a standard deviation of 50% of the mean, which is similar to what was observed in the tracing data. Rouse polymer models with and without loop extrusion were established using Brownian dynamics in the Euler scheme, as described in detail in Supplementary Note 2.

### Visualization and annotation of sequencing data

HiC maps and CHIP-seq data were visualized using pyGenomeTracks ^72^. CTCF motif orientation was annotated on the CTCF CHIP-seq data usingfimo v5.3.0 with the MA0139.1 motif and a p-value threshold of 0.01. The motif overlapping the CTCF Chip-seq peak with the lowest fimo p-value was used.

## Supplementary Materials

Supplementary Notes 1-2

Supplementary Figs. S1 to S8

Supplementary Data S1 to S2

## References

1. Dekker, J. & Mirny, L. The 3D Genome as Moderator of Chromosomal Communication. Cell 164, 1110–1121 2016.

2. Dixon, J. R. et al. Topological domains in mammalian genomes identified by analysis of chromatin interactions. Nature 485, 376–380 2012.

3. Rao, S. S. P. et al. A 3D Map of the Human Genome at Kilobase Resolution Reveals Principles of Chromatin Looping. Cell 159, 1665–1680 2014.

4. Rowley, M. J. & Corces, V. G. Organizational principles of 3D genome architecture. Nat Rev Genet 19, 789–800 2018.

5. Wutz, G. Topologically associating domains and chromatin loops depend on cohesin and are regulated by CTCF, WAPL, and PDS5 proteins. The EMBO Journal 36, 3573–3599 2017.

6. Rao, S. S. P. et al. Cohesin Loss Eliminates All Loop Domains. Cell vol. 171 https://pubmed.ncbi.nlm.nih.gov/28985562/ 2017.

7. Davidson, I. F. et al. DNA loop extrusion by human cohesin. Science 366, 1338–1345 2019.

8. Kim, Y., Shi, Z., Zhang, H., Finkelstein, I. J. & Yu, H. Human cohesin compacts DNA by loop extrusion. Science 366, 1345–1349 2019.

9. Fudenberg, G. et al. Formation of Chromosomal Domains by Loop Extrusion. Cell Reports 15, 2038–2049 2016.

10. Sanborn, A. L. et al. Chromatin extrusion explains key features of loop and domain formation in wild-type and engineered genomes. PNAS 112, E6456–E6465 2015.

11. Vian, L. et al. The Energetics and Physiological Impact of Cohesin Extrusion. Cell 173, 1165-1178.e20 2018.

12. Gabriele, M. et al. Dynamics of CTCF- and cohesin-mediated chromatin looping revealed by live-cell imaging. Science 376, 496–501 2022.

13. Sati, S. & Cavalli, G. Chromosome conformation capture technologies and their impact in understanding genome function. Chromosoma 126, 33–44 2017.

14. Maeshima, K., Ide, S. & Babokhov, M. Dynamic chromatin organization without the 30-nm fiber. Current Opinion in Cell Biology 58, 95–104 2019.

15. Beliveau, B. J. et al. Versatile design and synthesis platform for visualizing genomes with Oligopaint FISH probes. PNAS 109, 21301–21306 2012.

16. Beliveau, B. J. et al. Single-molecule super-resolution imaging of chromosomes and in situ haplotype visualization using Oligopaint FISH probes. Nature Communications 6, 2015.

17. Markaki, Y. et al. The potential of 3D-FISH and super-resolution structured illumination microscopy for studies of 3D nuclear architecture. BioEssays 34, 412–426 2012.

18. Boettiger, A. N. et al. Super-resolution imaging reveals distinct chromatin folding for different epigenetic states. Nature 529, 418–422 2016.

19. Wang, S. et al. Spatial organization of chromatin domains and compartments in single chromosomes. Science 353, 598–602 2016.

20. Bintu, B. et al. Super-resolution chromatin tracing reveals domains and cooperative interactions in single cells. Science 362, 2018.

21. Nir, G. et al. Walking along chromosomes with super-resolution imaging, contact maps, and integrative modeling. PLOS Genetics 14, e1007872 2018.

22. Cardozo Gizzi, A. M. et al. Microscopy-Based Chromosome Conformation Capture Enables Simultaneous Visualization of Genome Organization and Transcription in Intact Organisms. Molecular Cell 74, 212-222.e5 2019.

23. Mateo, L. J. et al. Visualizing DNA folding and RNA in embryos at single-cell resolution. Nature 568, 49–54 2019.

24. Su, J.-H., Zheng, P., Kinrot, S. S., Bintu, B. & Zhuang, X. Genome-Scale Imaging of the 3D Organization and Transcriptional Activity of Chromatin. Cell 182, 1641-1659.e26 2020.

25. Takei, Y. et al. Integrated spatial genomics reveals global architecture of single nuclei. Nature 590, 344–350 2021.

26. Solovei, I. et al. Spatial Preservation of Nuclear Chromatin Architecture during Three-Dimensional Fluorescence in Situ Hybridization (3D-FISH). Experimental Cell Research 276, 10–23 2002.

27. Robinett, C. C. et al. In vivo localization of DNA sequences and visualization of large-scale chromatin organization using lac operator/repressor recognition. J Cell Biol 135, 1685–1700 1996.

28. Barton, C. et al. ChromoTrace: Computational reconstruction of 3D chromosome configurations for super-resolution microscopy. PLOS Computational Biology 14, e1006002 2018.

29. Goodwin, E. & Meyne, J. Strand-specific FISH reveals orientation of chromosome 18 alphoid DNA. CGR 63, 126–127 1993.

30. Brown, J. M. et al. A tissue-specific self-interacting chromatin domain forms independently of enhancer-promoter interactions. Nature Communications 9, 3849 2018.

31. Brown, J. M., De Ornellas, S., Parisi, E., Schermelleh, L. & Buckle, V. J. RASER-FISH: non-denaturing fluorescence in situ hybridization for preservation of three-dimensional interphase chromatin structure. Nat Protoc 1–26 (2022) doi:10.1038/s41596-022-00685-8.

32. Schueder, F. et al. Universal Super-Resolution Multiplexing by DNA Exchange. Angewandte Chemie International Edition 56, 4052–4055 2017.

33. Luo, Y. et al. New developments on the Encyclopedia of DNA Elements (ENCODE) data portal. Nucleic Acids Res 48, D882–D889 2020.

34. Fudenberg, G. & Mirny, L. A. Higher order chromatin structure: bridging physics and biology. Curr Opin Genet Dev 22, 115–124 2012.

35. Landry, J. J. M. et al. The Genomic and Transcriptomic Landscape of a HeLa Cell Line. G3 Genes |Genomes| Genetics 3, 1213–1224 2013.

36. Karlsson, M. et al. A single–cell type transcriptomics map of human tissues. Science Advances 7, eabh2169 2021.

37. Alexanian, M. et al. A transcriptional switch governs fibroblast activation in heart disease. Nature 595, 438–443 2021.

38. Davidson, I. F. & Peters, J.-M. Genome folding through loop extrusion by SMC complexes. Nat Rev Mol Cell Biol 22, 445–464 2021.

39. Holzmann, J. et al. Absolute quantification of cohesin, CTCF and their regulators in human cells. eLife 8, e46269 2019.

40. Wutz, G. et al. ESCO1 and CTCF enable formation of long chromatin loops by protecting cohesinSTAG1 from WAPL. eLife 9, e52091 2020.

41. Hansen, A. S., Pustova, I., Cattoglio, C., Tjian, R. & Darzacq, X. CTCF and cohesin regulate chromatin loop stability with distinct dynamics. eLife 6, e25776 2017.

42. Tedeschi, A. et al. Wapl is an essential regulator of chromatin structure and chromosome segregation. Nature 501, 564–568 2013.

43. Casa, V. et al. Redundant and specific roles of cohesin STAG subunits in chromatin looping and transcriptional control. Genome Res. 30, 515–527 2020.

44. Nagano, T. et al. Single cell Hi-C reveals cell-to-cell variability in chromosome structure. Nature 502, 2013.

45. Mach, P. et al. Cohesin and CTCF control the dynamics of chromosome folding. Nat Genet 1–12 (2022) doi:10.1038/s41588-022-01232-7.

46. Darrow, E. M. et al. Deletion of DXZ4 on the human inactive X chromosome alters higher-order genome architecture. PNAS 113, E4504–E4512 2016.

47. Ran, F. A. et al. Double Nicking by RNA-Guided CRISPR Cas9 for Enhanced Genome Editing Specificity. Cell 154, 1380–1389 2013.

48. Yesbolatova, A., Natsume, T., Hayashi, K.-I. & Kanemaki, M. T. Generation of conditional auxin-inducible degron (AID) cells and tight control of degron-fused proteins using the degradation inhibitor auxinole. Methods 164–165, 73–80 2019.

49. Gans, J. D. & Wolinsky, M. Improved assay-dependent searching of nucleic acid sequence databases. Nucleic Acids Res 36, e74 2008.

50. Beliveau, B. J. et al. OligoMiner provides a rapid, flexible environment for the design of genome-scale oligonucleotide in situ hybridization probes. PNAS 115, E2183–E2192 2018.

51. Zadeh, J. N. et al. NUPACK: Analysis and design of nucleic acid systems. Journal of Computational Chemistry 32, 170–173 2011.

52. Georgakopoulos-Soares, I., Yizhar-Barnea, O., Mouratidis, I., Hemberg, M. & Ahituv, N. Absent from DNA and protein: genomic characterization of nullomers and nullpeptides across functional categories and evolution. Genome Biology 22, 245 2021.

53. Untergasser, A. et al. Primer3—new capabilities and interfaces. Nucleic Acids Research 40, e115 2012.

54. Xu, Q., Schlabach, M. R., Hannon, G. J. & Elledge, S. J. Design of 240,000 orthogonal 25mer DNA barcode probes. PNAS 106, 2289–2294 2009.

55. Moffitt, J. R. & Zhuang, X. Chapter One - RNA Imaging with Multiplexed Error-Robust Fluorescence In Situ Hybridization (MERFISH). in Methods in Enzymology (eds. Filonov, G. S. & Jaffrey, S. R.) vol. 572 1–49 (Academic Press, 2016).

56. Mateo, L. J., Sinnott-Armstrong, N. & Boettiger, A. N. Tracing DNA paths and RNA profiles in cultured cells and tissues with ORCA. Nat Protoc 16, 1647–1713 2021.

57. Xiang, W. et al. Correlative live and super-resolution imaging reveals the dynamic structure of replication domains. The Journal of cell biology 217, 1973–1984 2018.

58. Virtanen, P. et al. SciPy 1.0: fundamental algorithms for scientific computing in Python. Nature Methods 17, 261–272 2020.

59. Stringer, C., Wang, T., Michaelos, M. & Pachitariu, M. Cellpose: a generalist algorithm for cellular segmentation. Nature Methods 18, 100–106 2021.

60. Politi, A. Z. et al. Quantitative mapping of fluorescently tagged cellular proteins using FCS-calibrated four-dimensional imaging. Nature protocols 13, 1445–1464 2018.

61. The pandas development team. pandas-dev/pandas: Pandas. (2020) doi:10.5281/zenodo.3509134.

62. Walt, S. van der et al. scikit-image: image processing in Python. PeerJ 2, e453 2014.

63. Pedregosa, F. et al. Scikit-learn: Machine Learning in Python. Journal of Machine Learning Research 12, 2825–2830 2011.

64. Dask Development Team. Dask: Library for dynamic task scheduling. 2016.

65. napari contributors. napari: a multi-dimensional image viewer for python. 2019.

66. Czech, E., Aksoy, B. A., Aksoy, P. & Hammerbacher, J. Cytokit: a single-cell analysis toolkit for high dimensional fluorescent microscopy imaging. BMC Bioinformatics 20, 448 2019.

67. McInnes, L., Healy, J. & Melville, J. UMAP: Uniform Manifold Approximation and Projection for Dimension Reduction. arXiv:1802.03426 [cs, stat] 2020.

68. Waskom, M. L. seaborn: statistical data visualization. Journal of Open Source Software 6, 3021 2021.

69. Arun, K. S., Huang, T. S. & Blostein, S. D. Least-Squares Fitting of Two 3-D Point Sets. IEEE Trans. Pattern Anal. Mach. Intell. 9, 698–700 1987.

70. Ramachandran, P. & Varoquaux, G. Mayavi: a package for 3D visualization of scientific data. Comput. Sci. Eng. 13, 40–51 2011.

71. Atkins, J. E., Boman, E. G. & Hendrickson, B. A Spectral Algorithm for Seriation and the Consecutive Ones Problem. SIAM J. Comput. 28, 297–310 1998.

72. Lopez-Delisle, L. et al. pyGenomeTracks: reproducible plots for multivariate genomic datasets. Bioinformatics 37, 422–423 2021.

