## Supplementary Materials for "Nanoscale 3D DNA tracing in single human cells visualizes loop extrusion directly in situ"

**This PDF file includes:**

Supplementary Notes
Supplemental Figs. S1 to S8
Captions for Supplemental Data S1 to S2

**Other Supplementary Materials for this manuscript include the following:**

Data S1 to S2

### Supplementary Notes

#### *Supplemental Note 1: Validation of DNA FISH protocols for high resolution chromatin tracing*

To investigate which FISH protocol would best preserve chromatin structure and yet allow the high labelling efficiency required for chromatin tracing [1] we compared acid denaturing FISH at 86°C [2–6] or 75°C [7] with enzymatic, non-denaturing FISH (CO-FISH/RASER-FISH; see schematic in Supp. Fig S1a) [8–12]. We first labelled all genomic DNA of human diploid RPE-1 cells with DAPI and a 10 kb single-copy locus in the MYC gene on chromosome 8 with 96 oligopaint probes (Supp. Fig S1b) to compare overall genome structure preservation and FISH labelling efficiency. Diffraction limited imaging of cells after denaturing FISH at 86°C showed strong nuclear structure perturbations such as micrometer scale DNA leakages into the cytoplasm [13] and more than the two expected FISH signals from both alleles of the MYC gene. Lowering the temperature of denaturing FISH to 75°C decreased large scale DNA structure artefacts but also led to lower FISH-labelling efficiency (Supp. Fig. S1b, c). Non-denaturing FISH on the other hand showed very good overall DNA structure preservation and only slightly reduced FISH-labeling (Supp. Fig. S1b, c, d).

To investigate chromatin structure preservation at a smaller scale, we imaged cells with structured illumination microscopy (SIM) [7] in which TAD-sized co-replicating domains had been labelled with a pulse of fluorescent dUTPs during DNA replication [14]. While the initial fixation and permeabilization steps did not substantially alter genome or TAD level structure (Supp. Fig. S1e, f) at the scale of submicrometer domains, denaturation with hydrochloric acid and/or 86°C [3,5,6,15] led to strong structural perturbations (Supp. Fig. S1g, h, k, n), including fragmentation of heterochromatin, loss of nuclear integrity, spilling of DNA into the cytoplasm, and displacement/loss of co-replicating domains, consistent with previous reports [11,13]. Denaturation at 75°C showed milder but still significant perturbations of small scale chromatin structure (Supp. Fig. S1i, l, n), while non-denaturing FISH showed the best preservation (Supp. Fig. S1j, k, l, n).

To compare the FISH protocols for 3D reconstruction of genome folds, we carried out pilot tracing experiments using ten 5 kb probe-sets tiled along a 100 kb region upstream of the MYC gene. Here, denaturing FISH frequently showed fragmentation of one 5 kb locus into multiple spots suggesting chromatin disintegration, incompatible with high precision chromatin tracing (Supp. Fig. S1o). By contrast, non-denaturing FISH showed highly consistent labelling of single diffraction-limited spots for each of the ten 5 kb loci (Supp. Fig. S1p), which is the expected behaviour of native chromatin [14,16].

#### *Supplemental Note 2: Loop extrusion and Rouse polymer model formulation*

Chromatin was modelled with a Rouse polymer using Brownian dynamics similar to previous approaches [17,18]. The model was initially derived by Rouse from the Langevin motion equation in the over-damped regime [19]:

$$\frac{d}{dt}\boldsymbol{r_i} = \frac{\kappa}{\gamma}[(r_{i-1} - r_i) + (r_{i+1} - r_i)] + \sqrt{6D}\frac{d\xi}{dt}$$

Where $\mathbf{r_i}$ is the position of the ith bead, $\kappa$ is the spring constant of the Hookean spring coupling the beads, $\gamma$ is the friction coefficient, D is the diffusion coefficient of each bead, while $\xi$ is a Gaussian noise with zero mean and unit variance. The rouse polymer was simulated in an Euler scheme:

$$r_i(t + \Delta t) = r_i(t) + \frac{\kappa}{\gamma}[(r_{i-1} - r_i) + (r_{i+1} - r_i)] + \sqrt{6D\Delta t}\xi$$

with $\Delta t = 0.01s$, and each bead a monomer of 1 kb. There are numerous estimates of the diffusion coefficient of chromatin depending on experimental approaches, see e.g. [20–23]. We chose an intermediate value [23] for our simulation and swept $\frac{\kappa}{\gamma}$ values until reaching a polymer simulation showing physical distance scaling similar to high resolution tracing of 300 kb regions in cohesin-depleted cells ($r^2$= 0.996 for MEOX1 region). The values for all simulation parameters are listed in Supp. Table 1. In addition, a hard-shell potential set at 20 nm was used between interacting beads to avoid spatial overlap.

Loop extrusion was simulated by adding an extra spring between non-adjacent monomers corresponding to the left and right sides of a single stepping Cohesin, with the same spring constant as between adjacent monomers. Cohesin was modeled as two populations (Cohesin-STAG1 and Cohesin-STAG2), with their experimentally measured cellular abundances [24] distributed proportionally onto the simulated polymer, and off-rates corresponding to residence times matching kinetic FRAP data of STAG1 and STAG2 in CTCF-depleted cells [25]. On-rates were set so that the average bound fraction matched experimental data [24–26]. Upon binding, each Cohesin was placed at a random position on the polymer and while bound each side stepped left and right, respectively, along the polymer with a defined rate (extrusion rate) matching those measured by in vitro experiments [27,28] and estimated from modeling HiC data [29]. If one side stepped outside the polymer, the Cohesin was unbound.

The boundary factor CTCF was modelled with an off-rate from the residence time measured from FRAP and single particle tracking experiments [24]. Experimentally measured cellular abundances of CTCF [26] were distributed among all CTCF Chip-seq sites with annotated motif orientations (see methods). Upon binding, each CTCF would target one of the peak sites in the simulated region with a probability matching the relative signal of the Chip-seq peak in the simulated region. We note that this simplified approach of distributing CTCFs could lead to occasional discrepancy between the model and experimental data, as CTCF boundary strength and Chip-seq signal magnitude are not always directly correlated [30].

At each timestep, binding or unbinding of Cohesin and CTCF occurred with a probability of the inverse of the respective residence time, giving an exponentially distributed lifetime corresponding to the FRAP or single molecules measurements [24–26].

We made the following assumptions of the interactions between CTCF and Cohesin based on recent studies [25,31–33]: If a Cohesin encountered a CTCF in a convergent orientation, the respective side was stalled as long as the CTCF remained bound, and a new, extended lifetime in the CTCF-bound state was used [25,33]. CTCF residence time remained unchanged [26]. Each CTCF could bind one Cohesin, while each Cohesin could bind two CTCFs, one with each side, and remained stalled on the respective side until the corresponding CTCF unbound. No interactions between Cohesins were included in the model.

Loop extrusion simulations were performed by adding Cohesins and CTCFs in an unbound state and equilibrated with 2000 initiation loop-extrusion timesteps (1 second steps), before stochastically halting before 4000 simulation steps were reached. To reduce boundary effects, loop extrusion simulations were performed on a genomic region padded by 2 Mb on each side of the targeted genomic region. The resulting Cohesin loops were used to simulate a Rouse polymer.
Each Rouse polymer simulation was initialized from a random walk with a step size standard deviation of $\sqrt{6D}$ of a region padded with 100 kb on each side of the genomic target, and pre-equilibrated with 10 000 timesteps (0.01 s) before adding the Cohesin loops and running for another 10 000 steps. Each loop extrusion and corresponding polymer simulation was repeated to generate 3000 simulated traces. Gaussian noise giving rise to a median 50 nm 3D error was added to simulate conservative experimental conditions.
After calibration against tracing data from CTCF-depleted HeLa cells (see main text), all simulations were run using only the Cohesin-STAG1 population (see Supp. Fig. S7c). The Cohesin-STAG1 bound fraction depends on CTCF binding, increasing from ~0.5 to ~0.8 in the presence of CTCF [25]. In pilot simulations using the measured cellular abundances of CTCF (~1.7 CTCFs per Chip-seq site), a bound fraction of 0.70±0.06 was found for Cohesin-STAG1 from nine ~5 Mb genomic regions on different chromosomes. This bound fraction increased to 0.83±0.06, close to the experimental value, when an abundance of 3 CTCFs per Chip-seq site was used in the same regions. This value for total CTCF abundance was used for further simulations.
ΔWAPL simulations were performed by setting the Cohesin-STAG1 residence time to the same value as the CTCF-bound lifetime. 5000 initiation steps and up to 5000 running steps (average 2500) were used in the loop extrusion simulation to approximate a 2h WAPL depletion. All simulation code is available as part of the LoopTrace package (see Code availability).

**Supplementary Table 1: Simulation parameters used for loop extrusion simulations**

| Parameter | Value | Reference |
|---|---|---|
| D | 3.4e-3 µm^2/s | [23] |
| $\frac{\kappa}{\gamma}$ | 0.13 | Estimated from HeLa ΔRad21 measurements. |
| CTCFs per Chip-seq site | 1.7 (initial), 3 (adjusted) | Initial estimate from [24], adjusted to achieve Cohesin-STAG1 bound fraction of ~0.8. |
| Total Cohesin per Mb | 250 000/7900 Mb = 37 | [24] |
| Cohesin-STAG1/STAG2 ratio | 0.2 | [24] |
| Cohesin-STAG1 CTCF-free bound fraction | 0.5 | [25] |
| Cohesin-STAG1 bound fraction with CTCF | 0.8 | [25] |
| Cohesin-STAG1 chromatin bound lifetime | 900 s | [25] |
| Cohesin-STAG1 CTCF bound lifetime | 18 000 s | [25] |
| Cohesin-STAG1 unbound lifetime | 900 s | Estimated from bound fraction |
| Cohesin-STAG2 CTCF-free bound fraction | 0.45 | [25] |
| Cohesin-STAG2 bound fraction with CTCF | 0.5 | [25] |
| Cohesin-STAG2 chromatin bound lifetime | 480 s | [25] |
| Cohesin-STAG2 CTCF bound lifetime | 900 s | [25] |
| Cohesin-STAG2 unbound lifetime | 620 s | Estimated from bound fraction |
| Cohesin extrusion rate | 0.375 kb/s (0.1875 kb/s per side) | [27–29] |
| CTCF bound fraction | 0.5 | [26] |
| CTCF chromatin bound lifetime | 120 s | [26] |
| CTCF unbound lifetime | 120 s | [26] |

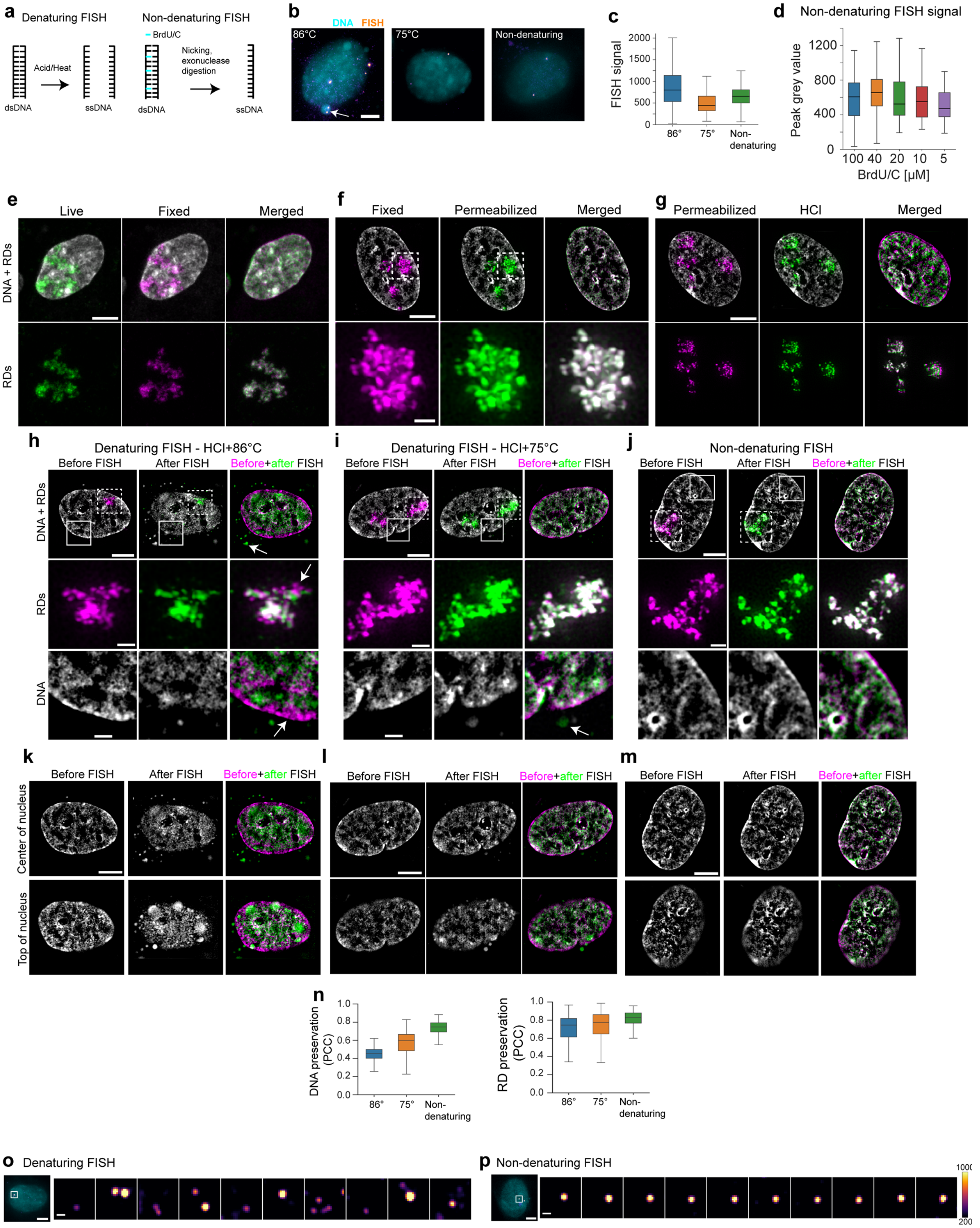
a Denaturing FISH
Non-denaturing FISH
BrdU/C
Acid/Heat
Nicking, exonuclease digestion
dsDNA
ssDNA
dsDNA
ssDNA
b DNA FISH
86°C
75°C
Non-denaturing
c FISH signal
86°
75°
Non-denaturing
d Non-denaturing FISH signal
Peak grey value
BrdU/C [μM]
e Live Fixed Merged
DNA + RDs
RDs
f Fixed Permeabilized Merged
g Permeabilized HCl Merged
h Denaturing FISH - HCl+86°C
Before FISH After FISH Before+after FISH
DNA + RDs
RDs
DNA
i Denaturing FISH - HCl+75°C
Before FISH After FISH Before+after FISH
j Non-denaturing FISH
Before FISH After FISH Before+after FISH
k Before FISH After FISH Before+after FISH
Center of nucleus
Top of nucleus
l Before FISH After FISH Before+after FISH
m Before FISH After FISH Before+after FISH
n DNA preservation (PCC)
RD preservation (PCC)
86°
75°
Non-denaturing
o Denaturing FISH
p Non-denaturing FISH

**Fig. S1.**

**a,** Illustration of the alternative FISH approaches investigated in this work. **b,** Diploid RPE-1 cells were prepared with alternative FISH protocols and labelled with 96 primary oligonucleotide FISH probes targeting a single copy 10 kb region upstream of the MYC gene on chromosome 8, visualized with an Atto565-labelled imaging oligo. FISH signal in extra-nuclear DNA material is indicated with the arrow. Maximum z-projections are shown. **c,** Peak signal intensity over background of FISH spots fit with a 3D Gaussian function for alternative FISH protocols. Data from n>50 cells per condition, data from one representative of two independent experiments. **d,** FISH signal with different total BrdU + BrdC concentrations (3:1 ratio) used for non-denaturing FISH. Data from n>50 cells per condition, data from one representative of two independent experiments. **e-g**, Comparison of preservation of chromatin structure during individual steps of the FISH protocol assessed by confocal imaging (e) or structured illumination microscopy (SIM, f-m) of cells labelled with Hoechst (DNA, grey) and Atto647-dUTP (replication domains, RDs, magenta/green).: **e,** from live to fixed cells; **f,** from fixed to permeabilized cells; and **g,** from permeabilization to acid treatment. Images representative of >10 (**e**) or >50 cells (**f, g**) in two independent experiments. **h-j,** Structural preservation of nuclear architecture after alternative FISH protocols, Extra-nuclear DNA material and substantial shifts in RD position indicated by arrows. Single z-planes are shown. **k-m,** Comparison of central and apical planes of cell nuclei before and after FISH treatment for the conditions described in **h-i**. **l,** Pearson's correlation of signal intensity of 3D drift-corrected SIM images before and after FISH. **h-l,** Data from n > 350 cells per condition from two independent experiments. **m-n,** Comparison of 3D sequential FISH imaging of probes targeting ten 5 kb regions tiled across 100 kb in the MYC locus using **e**, high-temperature (86°C) denaturing FISH and **f**, non-denaturing FISH. Single z-slices are shown. Data representative of **m,** 431 cells in one experiment or **n,** 562 cell in 3 experiments. Scale bars, 5 µm; RDs in **f; h-j**, 1 µm.

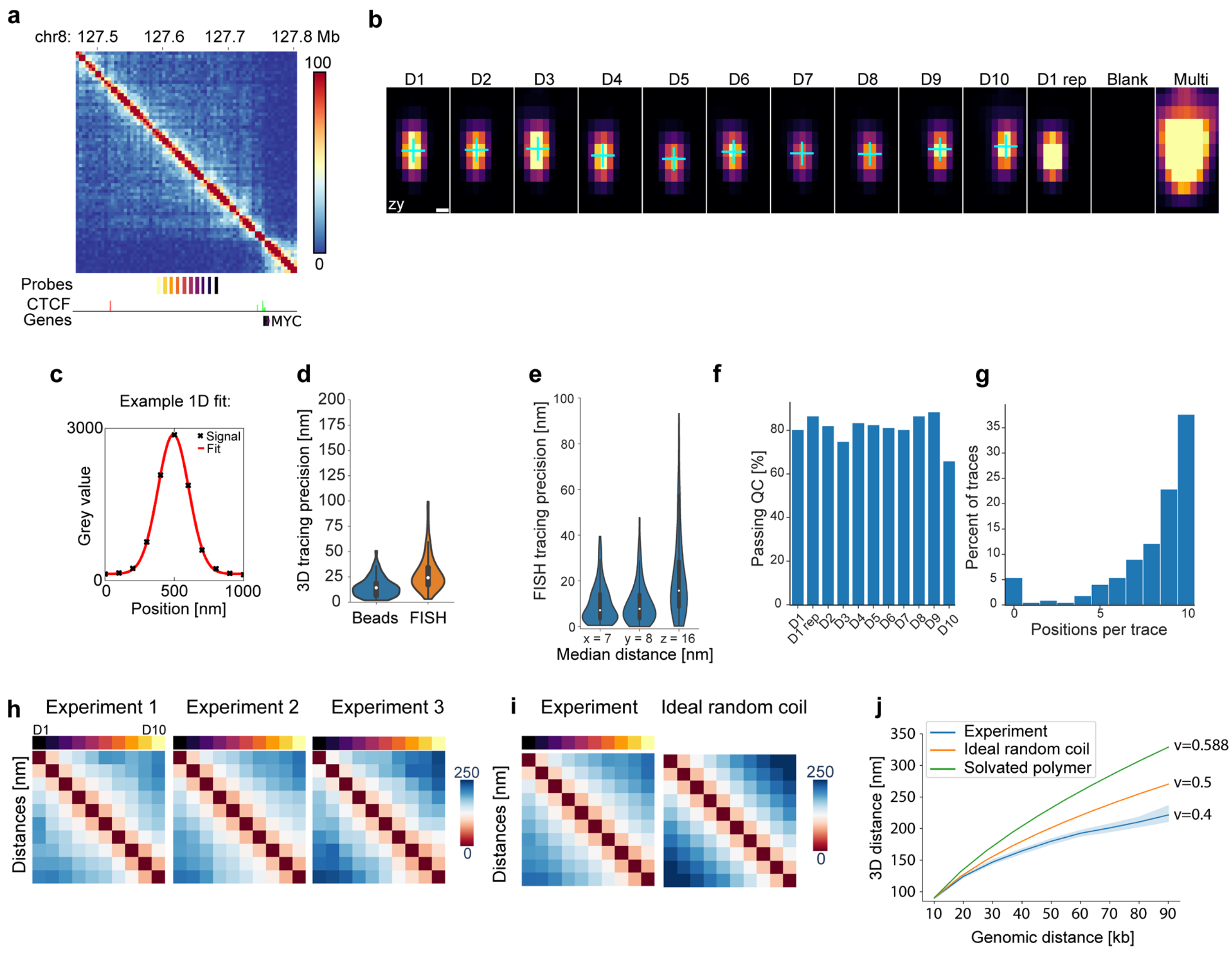


**Fig. S2.**

**a,** Hi-C contact map of RPE-1 cells (GSE71831) at 10 kb binning of the traced region in the vicinity of the MYC gene, with probe positions, CTCF chip-seq peaks indicated by direction of the predicted overlapping binding motif (green forward, red reverse), and genes (GENCODE v41). The probes cover 10 regions of 5 kb with 5-kb-gaps, spanning a total of 100 kb. **b**, Individual FISH signals from slices along the zy axis at the central x position of each fit from sequential hybridization of the 10 labelled regions (D1-D10), as well as repeat labelling (D1 rep), wash buffer only (Blank) or simultaneous labelling with multiple probes for region identification (Multi). Fit centroid positions are indicated by blue crosses. **c**, Example of 1D-signal (black) from a FISH spot, and a 1D-Gaussian fit (red line) to the signal that is used for centroid fitting. Tracing precision judged by **d**, 3D-Euclidean distance and **e**, individually in x, y and z-directions, across 10 hybridizations. Measurements made on fiducial beads (n=66, median=14 nm) or by re-hybridizing the first position after tracing (n=161 traces, median=24 nm). Data from one representative of 8 experiments. **f,** The percentage of spots detected and passing quality control (QC) for each hybridization in all traces. **g,** Percentage of traces with indicated number of hybridizations detected and passing QC. 80% of sequential hybridizations were detected in the majority of imaged cells.

Data in (**d-g)** from 223 traces in one representative experiment. **h,** Distance maps for tree replicate experiments tracing the 100 kb region upstream of the MYC gene on chromosome 8. Experimental replicates were highly consistent with average Spearman's r = 0.99. n=138, 203 and 182 traces for experiments 1, 2, and 3, respectively. Scale bar in nm. **i,** Average distance maps for experimental data and an ideal random coil model. **j,** Scaling relations between genomic and physical distance (median±95% c.i. estimated by bootstrapping) for the 100 kb region shown in (**h).**

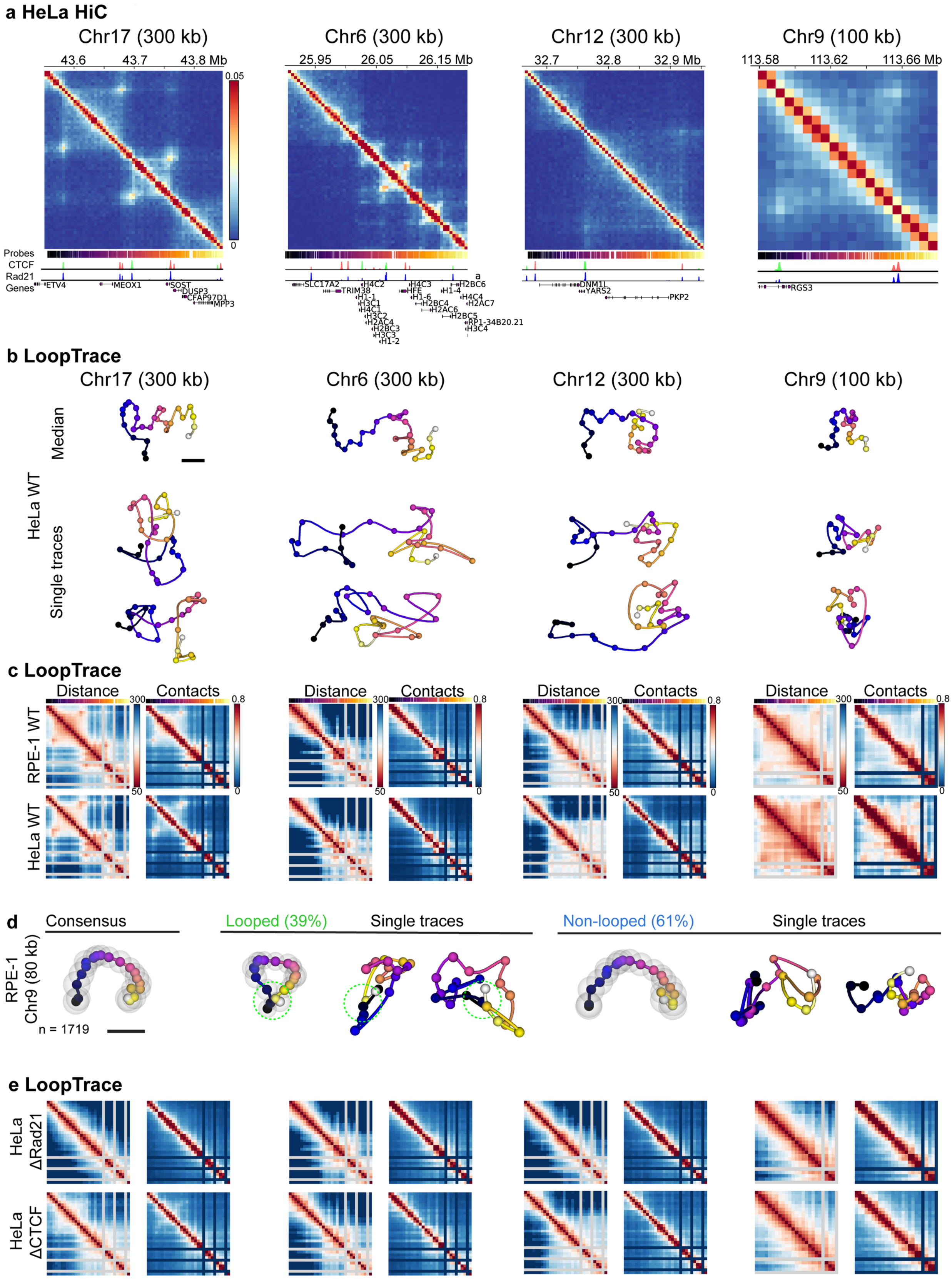
a HeLa HiC
Chr17 (300 kb)
Chr6 (300 kb)
Chr12 (300 kb)
Chr9 (100 kb)
43.6
43.7
43.8 Mb
25.95
26.05
26.15 Mb
32.7
32.8
32.9 Mb
113.58
113.62
113.66 Mb
0.05
0
Probes
CTCF
Rad21
Genes
ETV4
MEOX1
SOST
DUSP3
CFAP97D1
MPP3
SLC17A2
TRIM38
HFE
DNM1L
YARS2
PKP2
RGS3
b LoopTrace
Median
HeLa WT
Single traces
c LoopTrace
Distance
Contacts
300
50
0.8
0
RPE-1 WT
HeLa WT
d
Consensus
Looped (39%)
Single traces
Non-looped (61%)
RPE-1
Chr9 (80 kb)
n = 1719
e LoopTrace
HeLa
ΔRad21
HeLa
ΔCTCF

**Fig. S3.**

**a,** Deep HeLa HiC maps (from the 4DN data portal, accession number 4DNFIBM9QCFG) binned at 5 kb, as well as probe positions, CTCF and Rad21 Chip-seq (from the ENCODE portal) and gene annotations of 100-300 kb regions on chromosomes 17, 6, 12 and 9 that were targeted with probes for tracing. **b,** Trace reconstructions from median pairwise distances and representative single cell example traces from the indicated regions traced at 10 kb resolution across 300 kb, or 4 kb resolution across 100 kb in RPE-1 and HeLa Kyoto wild type (WT) cells. Experimental data from two independent experiments per condition with the indicated number of traces. **c,** Pairwise distance maps and contact frequencies (<120 nm) of tracing data from regions shown in (**b**)**.** Poorly performing secondary imager probes were filtered from the data (grey/dark blue stripes). **d,** Consensus traces, as well as looping and non-looping subsets of traces and representative single trace examples from the 80 kb looping region subset from the 100 kb region traced on chromosome 9 in RPE-1 cells. Proximity (<120 nm) between the positions of the looping CTCF-sites is indicated by dashed circles. **e,** Pairwise distance maps and contact frequencies of the same regions as shown in (**c**) from HeLa Kyoto Rad21-mEGFP-AID or CTCF-mEGFP-AID cells acutely depleted of Rad21 or CTCF by 2 h auxin treatment. Experimental data from two independent experiments per condition with the indicated number of traces. Spearman correlations between experimental conditions are available in Supplementary Data S2.

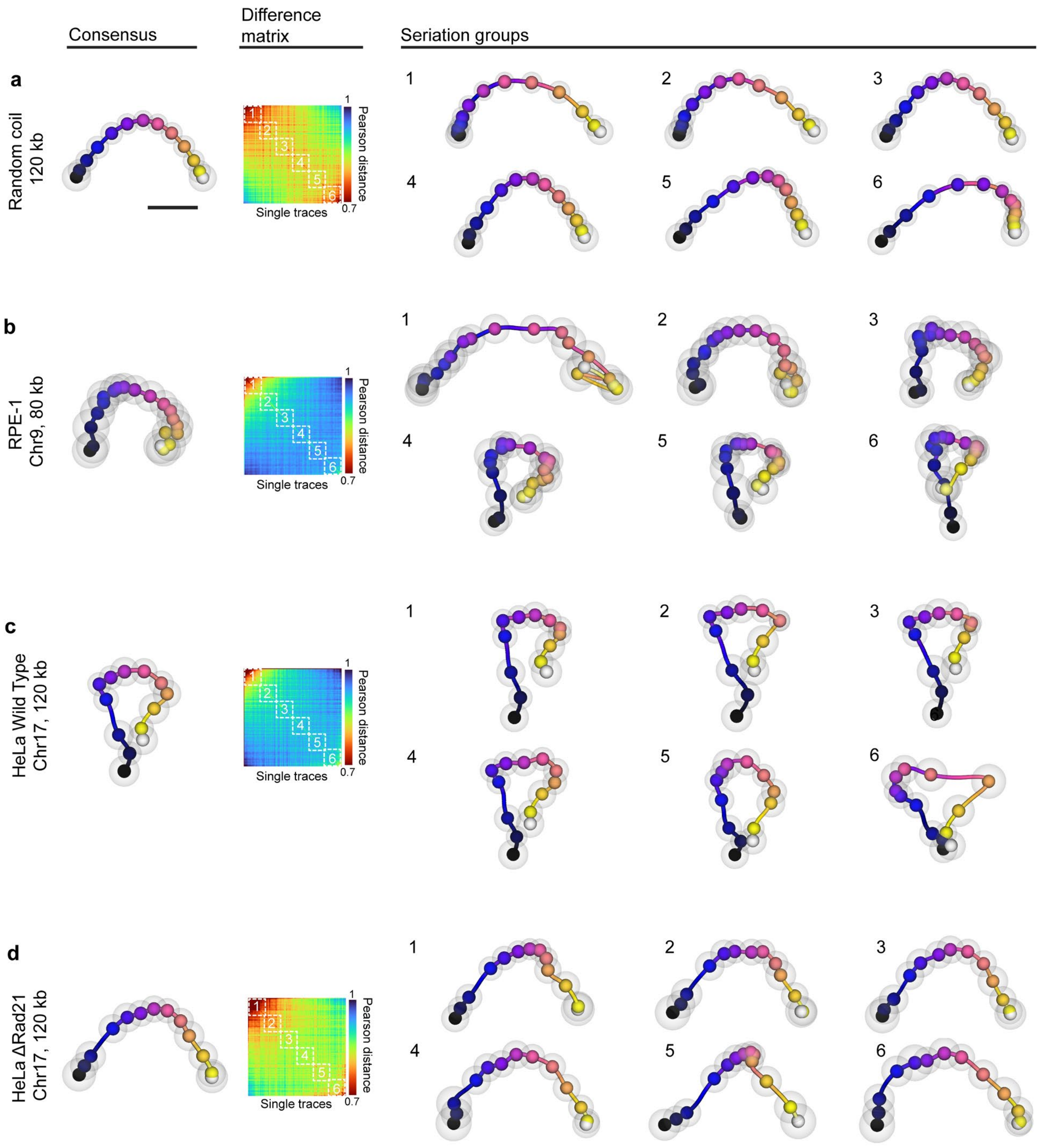


**Fig. S4.**

Consensus traces, difference matrices (1-$\sqrt{PCC}$ between all traces) sorted by their Fiedler vector, and consensus traces from seriation groups containing traces indicated with dashed squares on the difference matrix from **a**, simulated 120 kb ideal random coils, n=3000; **b,** RPE-1 cells from a 80 kb region (Chr9: 113.56-113.64 Mb), n=1719, 2 replicates. **c,** HeLa WT cells from a 120 kb region (Chr17: 43.5-43.6 Mb), n=2807, 2 replicates; **d,** HeLa ΔRad21 cells from a 120 kb region (Chr17: 43.5-43.6 Mb), n=2199, 2 replicates;

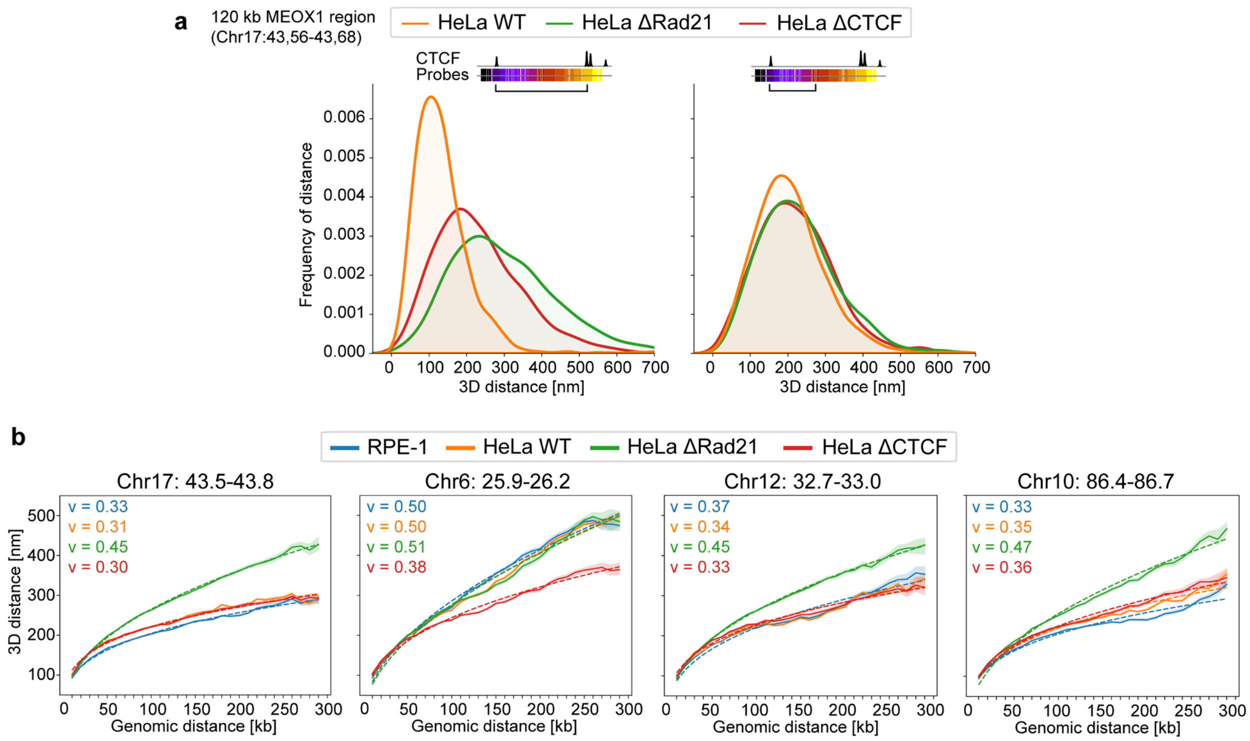


**Fig. S5.**

**a**, Distribution of 3D distances between the indicated probe positions in the traced region near the MEOX1 gene on Chr17 in HeLa WT cells and HeLa AID cells acutely depleted of Rad21 or CTCF by 2 h auxin treatment. **b,** Distance scaling relations from four 300 kb regions traced in the indicated cell lines (median±95% c.i. estimated by bootstrapping), with power law fits shown as dashed lines and resulting exponents as indicated. All fit parameters are listed in Supplementary Data S2. Data in (**a**) and (**b**) from 1000-3000 traces and two replicates per condition.

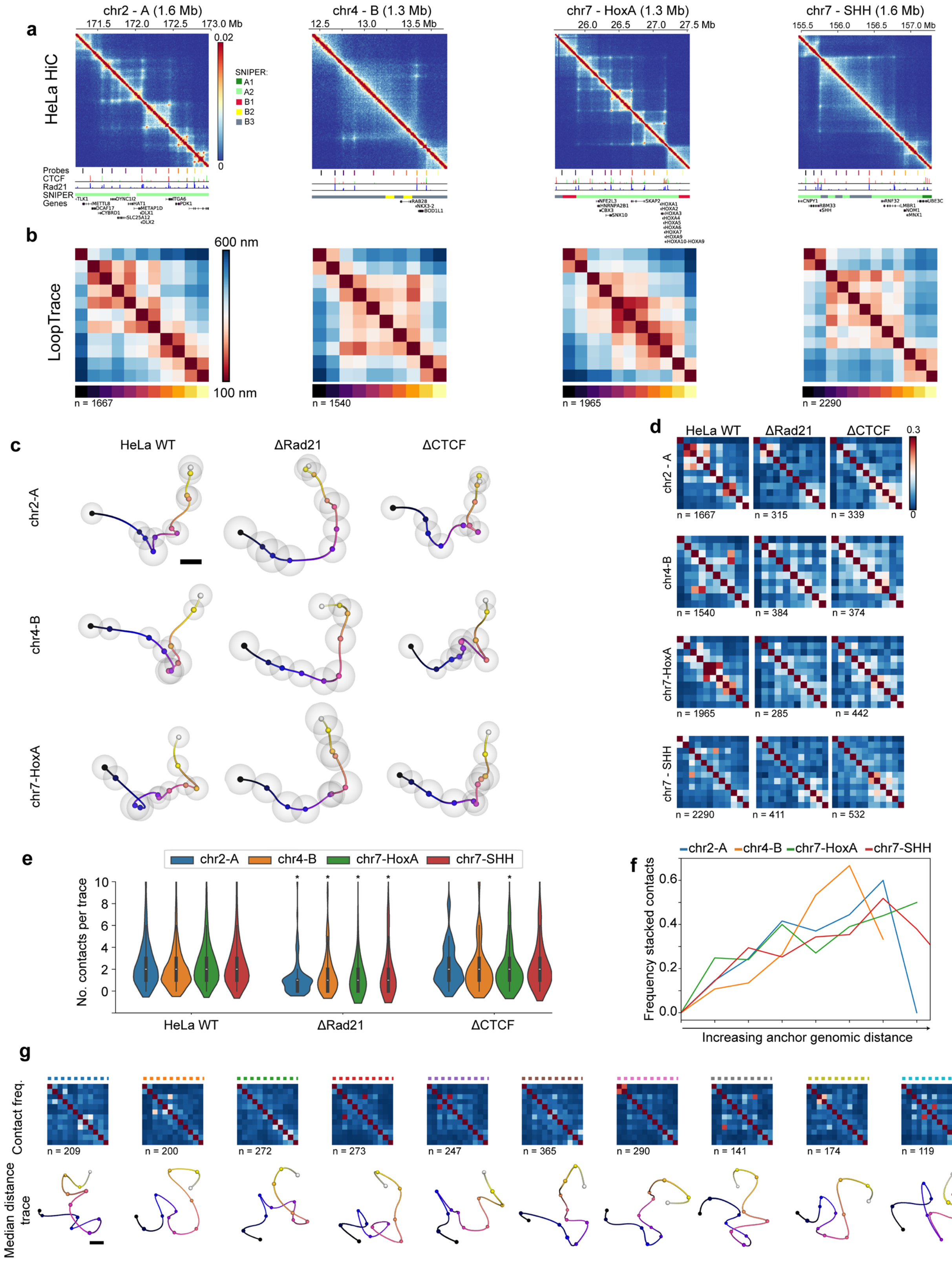
a
HeLa HiC
chr2 - A (1.6 Mb)
171.5
172.0
172.5
173.0 Mb
0.02
0
SNIPER:
A1
A2
B1
B2
B3
Probes
CTCF
Rad21
SNIPER
Genes
chr4 - B (1.3 Mb)
12.5
13.0
13.5 Mb
chr7 - HoxA (1.3 Mb)
26.0
26.5
27.0
27.5 Mb
chr7 - SHH (1.6 Mb)
155.5
156.0
156.5
157.0 Mb
b
LoopTrace
600 nm
100 nm
n = 1667
n = 1540
n = 1965
n = 2290
c
HeLa WT
ΔRad21
ΔCTCF
chr2-A
chr4-B
chr7-HoxA
d
HeLa WT
ΔRad21
ΔCTCF
0.3
0
chr2 - A
n = 1667
n = 315
n = 339
chr4-B
n = 1540
n = 384
n = 374
chr7-HoxA
n = 1965
n = 285
n = 442
chr7 - SHH
n = 2290
n = 411
n = 532
e
chr2-A
chr4-B
chr7-HoxA
chr7-SHH
No. contacts per trace
HeLa WT
ΔRad21
ΔCTCF
f
Frequency stacked contacts
Increasing anchor genomic distance
g
Contact freq.
1.0
0
n = 209
n = 200
n = 272
n = 273
n = 247
n = 365
n = 290
n = 141
n = 174
n = 119
Median distance trace

**Fig. S6.**

**a,** HeLa HiC maps (4DNFIBM9QCFG) binned at 10 kb of the four TAD-scale regions selected for tracing. The probes, as well as HeLa CTCF ChIP-seq peaks indicating motif directionality, HeLa Rad21 ChIP-seq peaks and SNIPER compartmental annotations are listed. **b,** Median 3D pairwise distance maps of measured probe positions in the four indicated regions in HeLa cells. Trace numbers as indicated from three experiments. **c,** Consensus traces from wild type HeLa Kyoto cells or HeLa Rad21-mEGFP-AID or CTCF-mEGFP-AID cells treated with auxin for 2 h. **d**, Contact frequencies (<120 nm) of the regions and treatment conditions in (**c**)**. e,** Number of contacts (<120 nm) per trace for the different regions and perturbations. * $p<0.05$ for condition vs wild type, tested by Kruskal-Wallis non-parametric 1-way ANOVA followed by Conover's test with Holm adjustment for multiple comparisons. **f,** Frequency of longer-range contacts established by stacking of shorter contacts sorted according to distance spanned by the largest contact. **g**, Contact frequencies and median distance reconstructions of the clusters in Fig. 3c indicated by coloured dashed lines**.** Data from three (wild type) or two (AID lines) experiments, number of traces per group as indicated. Scale bars 100 nm.

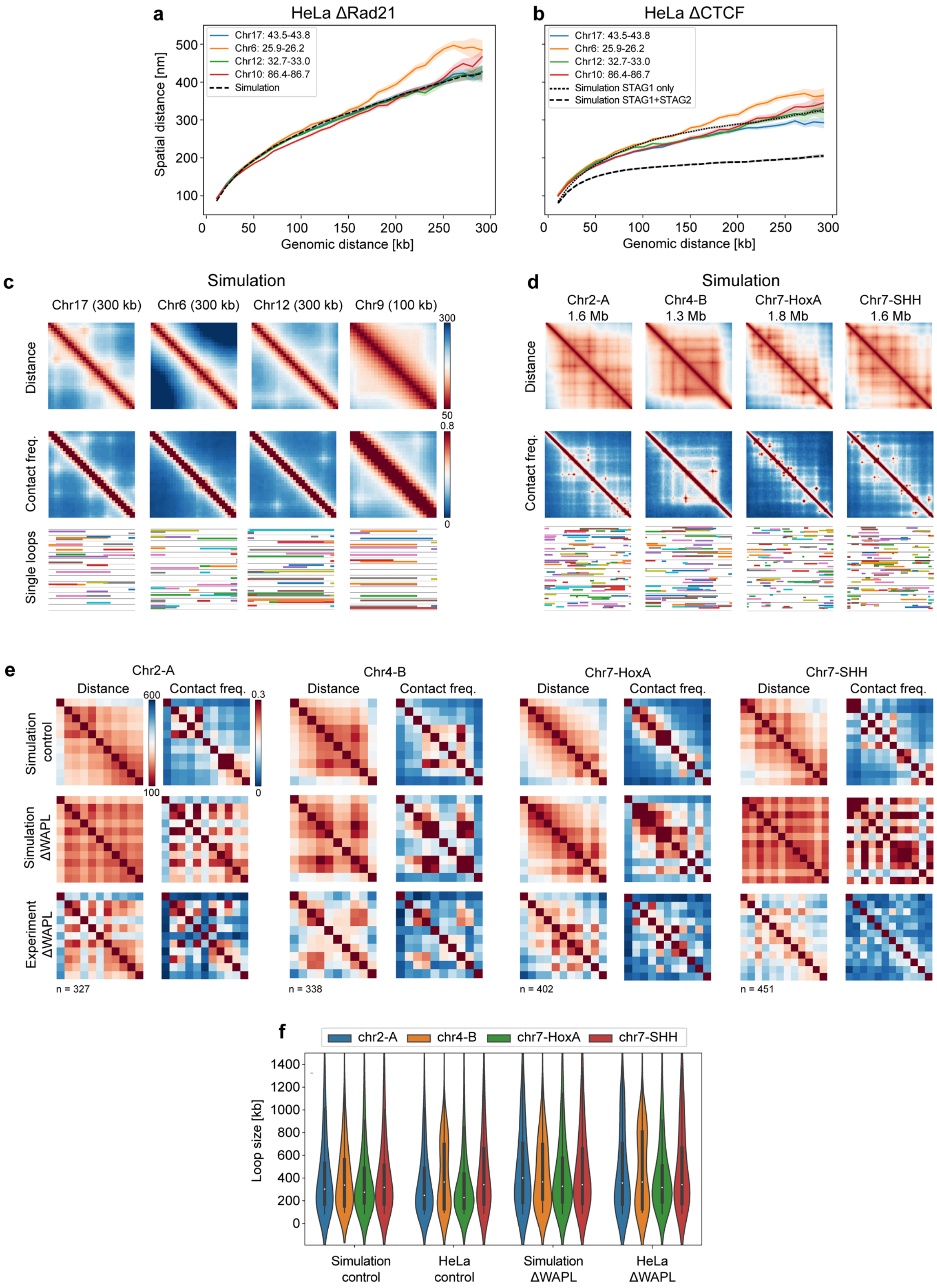
a
HeLa ΔRad21
Chr17: 43.5-43.8
Chr6: 25.9-26.2
Chr12: 32.7-33.0
Chr10: 86.4-86.7
Simulation
Spatial distance [nm]
Genomic distance [kb]
b
HeLa ΔCTCF
Simulation STAG1 only
Simulation STAG1+STAG2
c
Simulation
Chr17 (300 kb)
Chr6 (300 kb)
Chr12 (300 kb)
Chr9 (100 kb)
Distance
Contact freq.
Single loops
d
Chr2-A 1.6 Mb
Chr4-B 1.3 Mb
Chr7-HoxA 1.8 Mb
Chr7-SHH 1.6 Mb
e
Simulation control
Simulation ΔWAPL
Experiment ΔWAPL
n = 327
n = 338
n = 402
n = 451
f
chr2-A
chr4-B
chr7-HoxA
chr7-SHH
Loop size [kb]
Simulation control
HeLa control
Simulation ΔWAPL
HeLa ΔWAPL

**Fig. S7.**

**a,** Distance measurements from four 300 kb regions traced at 10 kb resolution in HeLa Kyoto Rad21-mEGFP-AID cells acutely depleted of Cohesin and corresponding calibrated Rouse polymer simulation with no loop extruders. **b,** Distance measurements from four 300 kb regions traced at 10 kb resolution in HeLa Kyoto CTCF-mEGFP-AID cells acutely depleted of CTCF and a corresponding Rouse polymer simulation with loop extruder abundances and dynamic parameters corresponding to Cohesin-STAG1+STAG2 or Cohesin-STAG1 only. **c,** Simulated data corresponding to the three 300 kb regions and one 100 kb region measured experimentally in Supp. Fig. 3. Examples of the ground truth loops underlying the simulations are displayed as colored lines spanning the genomic positions of the loops. **d,** Simulated data and exemplary ground truth loops corresponding to the four 1.3-1.8 Mb regions measured experimentally in Supp. Fig. 4. **e,** Experimental measurements from HeLa WAPL-Halo-AID cells treated for 2h by auxin and corresponding simulations of WAPL depletion (20-fold increased Cohesin residence time) from the four 1.3-1.8 Mb regions measured in Fig. 3. **f,** Distributions of loops lengths (excluding loops formed by stacking smaller loops) from the simulations or experimental measurements from WT and WAPL depletion conditions. The simulated data was sampled at the experimental probe positions and included corresponding probe drop-out rates as experiments for comparison. 3000 traces were simulated, number of traces for experimental conditions from two experimental replicates are indicated.

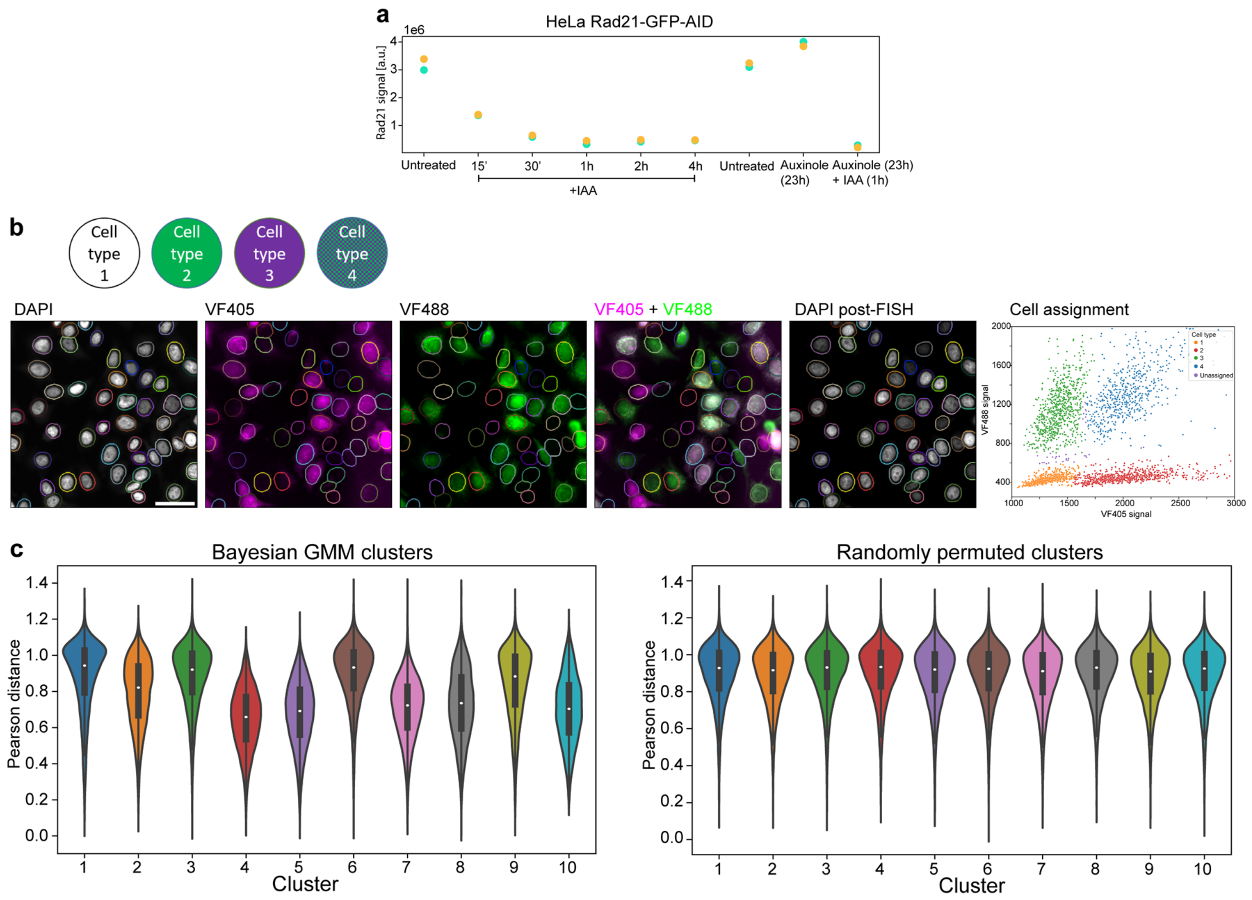


**Fig. S8.**

**a,** Chemiluminescence signal from simple Western assay using anti-Rad21 in untreated HeLa Rad21-mEGFP-AID cells or treated with auxin (IAA), auxinole or both for the indicated durations. Yellow and cyan markers indicate technical duplicates from one experiment. **b,** Overview of barcoding strategy to simultaneously acquire data from multiple cell lines. Each of up to four cell lines is either left unlabelled, or pre-labelled with VF405, VF488 or both before mixing and seeding the cells in the sample chamber. After fixation, VF405 and VF488 signals in cells in the entire sample chamber are imaged at lower magnification, then labelled with DAPI and reimaged before being processed for FISH. The same cells are relocated by registering the DAPI channel acquired during sequential FISH imaging with the pre-FISH images, and detecting the VF405 and VF488 intensities in a dilated nuclear mask (pseudo-coloured ellipses). The original cell types are then recovered by a manual gating strategy based on the VF405+VF488 intensity. Imaging representative of 100 fields of view in two experiments. **c,** All pairwise intra-cluster differences (using √(1-PCC) as distance metric) of traces from the Chr7-SHH region assigned to clusters shown in Fig. 3c and Supp. Fig. S6g (left), compared with randomly permuted cluster memberships (right). Overall differences detected by comparing the means of the real and permuted clusters ($p=0.001$, two-tailed t-test, total of $n=2290$ traces in 10 clusters). Results were identical for repeated permutations.

**Data S1. (separate file)**

All FISH probe oligonucleotide sequences used in this work.

**Data S2. (separate file)**

Spearman correlations and $r^2$ values between experimental conditions and simulated tracing data, and fit parameters from power law fits to distance measurements in tracing data.
